# Ginsentide TP1 Protects Hypoxia-Induced Dysfunction and ER Stress-Linked Apoptosis

**DOI:** 10.1101/2023.04.12.536670

**Authors:** Bamaprasad Dutta, Shining Loo, Antony Kam, Siu Kwan Sze, James P. Tam

## Abstract

Hypoxia-induced vascular endothelial dysfunction (VED) is a significant contributor to several severe human conditions, including heart disease, stroke, dementia, and cancer. However, current treatment options for VED are limited due to a lack of understanding of the underlying disease mechanisms and therapeutic leads. We recently discovered a heat-stable microprotein in ginseng, known as ginsentide TP1 that has been shown to reduce vascular dysfunction in cardiovascular disease models. In this study, we use a combination of functional assays and quantitative pulsed SILAC proteomics to determine new proteins synthesized in hypoxia and to show that ginsentide TP1 provides protection for human endothelial cells against hypoxia and ER stress. We found that hypoxia activates various pathways related to endothelium activation and monocyte adhesion, which in turn, impairs nitric oxide (NO) synthase activity, reduces NO bioavailability, and increases the production of reactive oxygen species that contribute to VED. Additionally, hypoxia triggers endoplasmic reticulum stress and initiates apoptotic signaling pathways associated with cardiovascular pathology. Treatment with ginsentide TP1 reduced surface adhesion molecule expression prevented activation of the endothelium and leukocyte adhesion, restored protein hemostasis, and reduced ER stress to protect against hypoxia-induced cell death. Ginsentide TP1 also restored NO signaling and bioavailability, reduced oxidative stress, and protected endothelial cells from endothelium dysfunction. In conclusion, this study shows that the molecular pathogenesis of VED induced by hypoxia can be mitigated by treatment with ginsentide TP1, which could be one of the key bioactive compounds responsible for the “cure-all” effect of ginseng. This research may lead to the development of new therapies for cardiovascular disorders.

## Introduction

Hypoxia-induced vascular endothelial dysfunction (VED) plays a key role in the pathogenesis of multiple human disorders, including heart disease, stroke, dementia, and diabetic complications; however, the molecular mediators of VED largely remain unknown. Clinical data suggest that hypoxia-induced by sleep apnea or high-altitude conditions is strongly associated with VED and may increase the risk of cardiovascular disorders and heart failure (1–6). Models of pulmonary vasoconstriction studied under low-oxygen conditions also suggest that hypoxia strongly influences the endothelium via connexin 40-mediated effects on endothelial membrane potential (7). However, hypoxia also modulates the functions of P2X and P2Y cell surface receptors, which stimulate the release of nitric oxide (NO) and hyperpolarizing factors (8). In addition, VED restricts blood flow and oxygen supply to tissues and endothelium, forming a vicious cycle and amplifying the hypoxia-induced VED. Acting through these underlying mechanisms, hypoxia can trigger proinflammatory endothelial responses, including the upregulation of cell adhesion molecules that increase macrophage infiltration (9), which is a known risk factor for adverse cardiovascular events (1, 4–6). Therapeutic targeting hypoxia-VED will break this vicious cycle and limit the progression of vascular pathology underlying major human diseases.

Under normal physiological conditions, NO generation maintains the quiescent state of the endothelium (10). In contrast, disrupting NO signaling triggers endothelial activation and the expression of cytokines, chemokines, and adhesion molecules that promote leukocyte adhesion and inflammation (10, 11). During oxidative stress, NO reacts with superoxide anions to generate peroxynitrite, which reduces the function of endothelial nitric oxide synthases (eNOSs) and limits NO production to promote VED (10, 12, 13). Hypoxia also induces endoplasmic reticulum (ER) stress, which triggers proapoptotic signaling pathways that are implicated in cardiovascular disease (14, 15). In contrast, the inhibition of ER stress-induced endothelial apoptosis is known to prevent VED and reduce the cardiovascular complications of diabetes (16). Therefore, ER stress represents a critical component of VED and a priority target for therapeutic interventions; however, methods for limiting ER stress in the hypoxic endothelium have not been identified.

Ginseng plants belonged to the *Panax* (panacea) genus and were named for their “cure-all” medicinal effects. For many centuries, ginseng has been used as an herbal medicine to treat various disorders, including heart disease and diabetes (17, 18). Currently, identified bioactive compounds are primarily small-molecule metabolites. Moreover, the molecular basis underlying the beneficial effects of ginseng on VED in human patients has not been clarified. Studies conducted in our laboratory led to the identification of a novel family of cysteine-rich peptides (CRPs) or microproteins from ginseng plants designated as ginsentides (19). To date, 14 ginsentides (TP1–14) have been identified from three common and medicinally valuable ginseng species, i.e., *Panax ginseng*, *Panax notoginseng*, and *Panax quinquefolius*. (19). All ginsentides are 31–33 amino acids long and contain a conserved 8-cysteine motif and 4 disulfide linkages (19). The prototypic member, ginsentide TP1 (hereafter given as TP1), is the most abundant CRP found in *P. ginseng* and *P. notoginseng*. The 31-residue TP1 is both cysteine- and glycine-rich (**Fig. 1A**). The eight cysteine residues in TP1 form a novel disulfide connectivity of Cys I–IV, II–VI, III–VII, and V–VIII, resulting in a pseudocyclic knot in which the first and last residues are cystine forming disulfides with a knotted and multicyclic structure (**Fig. 1A, B**) (19). This unique disulfide connectivity provides the structural rigidity to TP1 and braces it into a hyperstable and compact microprotein that is resistant to heat, acid, and both exopeptidase and endopeptidase degradation (19–22).

TP1 and other members of ginsentides (TP2-14) form a new CRP family. They contain eight cysteine residues and a cysteine motif similar to heveins found in *Hevea brasiliensis* and hevein-like peptides, but they can be distinguished from heveins by distinctly different disulfide connectivity, biosynthesis, and the absence of a chitin-binding domain which is essential for the host-defense functions of heveins and hevein-like peptides (19–24). Because ginsentide TP1 is a prototype, its biological functions have not been explored. Recently, we showed that ginsentides TP1, TP3, and TP8 modulate multiple biological systems to exert potent effects on vascular biology (25).

Here, we show that TP1 protects endothelial cells (ECs) from hypoxia-induced and stress-linked dysfunctions underlying VED pathogens by synthesizing new proteins critical to their adaptive responses, we used pulsed SILAC (pSILAC) quantitative proteomics to pulse/chase *de novo* protein synthesis in ECs subjected to low-oxygen stress with or without TP1. Guided by the pSILAC results, we further investigated how TP1 protects ECs from hypoxia stress using functional assays. Using these approaches, we show that TP1 protects human ECs from hypoxia-induced VED by preventing ER stress-mediated apoptotic signaling and by increasing cell survival. We also compared TP1 with ginsenosides, a family of well-studied bioactive metabolites from ginseng. Our findings showed that ginsenosides did not exert protective effects against hypoxia-induced VED when compared to the novel ginsentides.

**Fig. 1:**
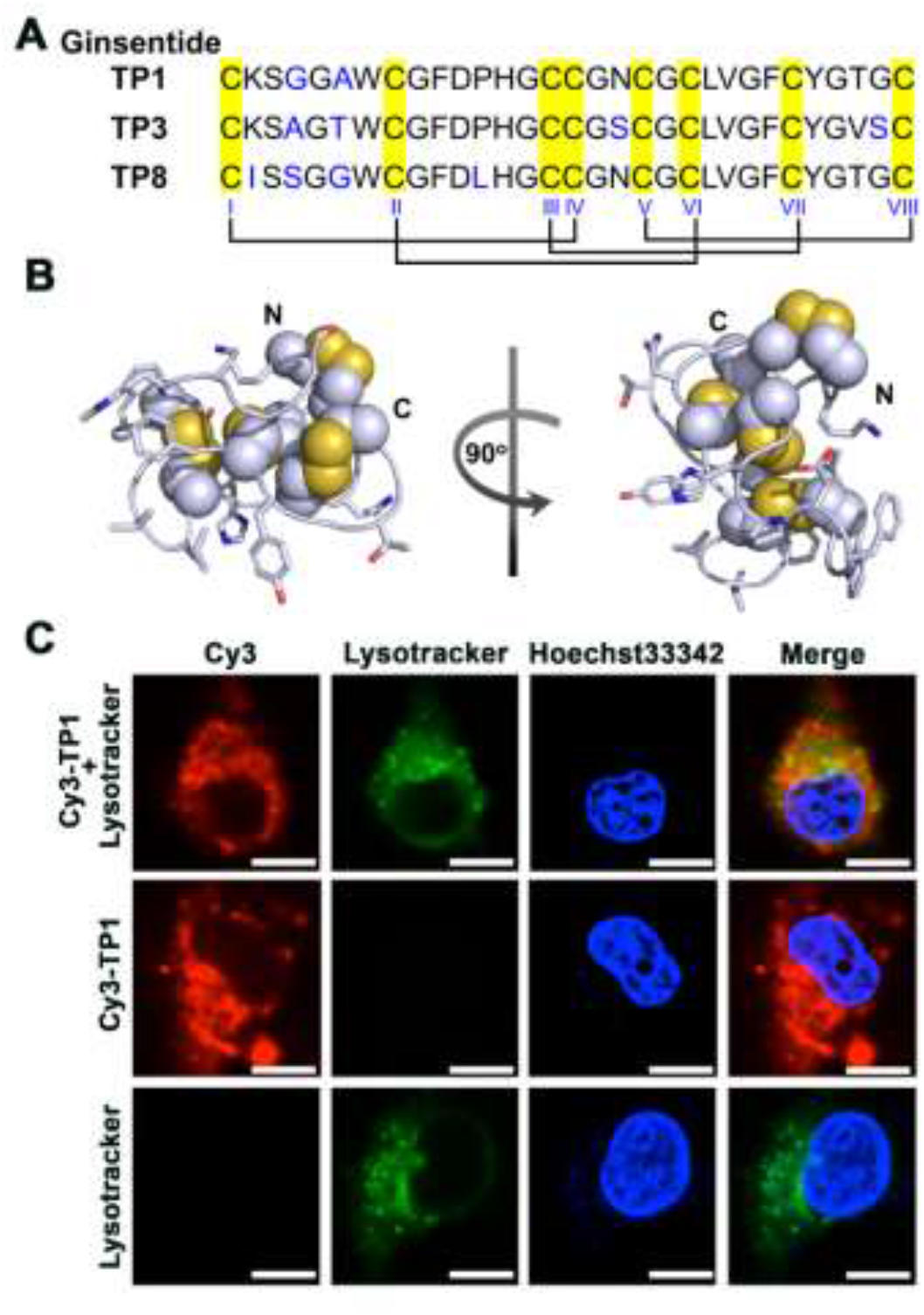
Ginsentides: cysteine- and glycine-rich cell-penetrating peptides from ginseng. A) Primary sequence and disulfide connectivity (Cys I–IV, Cys II–VI, Cys III– VII, and Cys V–VIII) of ginsentides TP1, TP3, and TP8. B) Cartoon representation of the NMR structure of the pseudocyclic TP1 (PDB code: 2ML7). The unusual disulfide connectivity between cysteine residues forms a pseudocyclic knot, in which the first and last residues are cysteines. C) Images of HUVEC-CS cells after incubation with 1 μM Cy3–TP1, captured using live-cell confocal microscopy at 37°C. Cy3–TP1 is shown in red, whereas lysosomes and nuclei are counterstained with LysoTracker (green) and Hoechst 33342 (blue), respectively. Scale bar: 5 µm.

## Results

### TP1 is a hyperstable nontoxic, and non-membranolytic cell-penetrating peptide

TP1 is a positively charged and hydrophobic peptide. To determine whether it interacts with intracellular proteins to protect against hypoxia-induced dysfunction, we site-specifically labeled TP1 with Cy3 at its lysine side chain. The cellular uptake of Cy3-labeled TP1 (Cy3–TP1) in live cells was observed using confocal microscopy to minimize artifacts in localizing labeled peptides under fluorescence microscopy (26). The live-cell images of human umbilical vein/vascular endothelium (HUVEC-CS) cells after incubation with 1 μM Cy3–TP1 for 1 h revealed that Cy3–TP1 was internalized and distributed throughout the cytoplasm, whereas LysoTracker counterstaining showed that Cy3–TP1 also accumulated in lysosomes (**Fig. 1C**).

To investigate the toxicity of TP1, we used a lactate dehydrogenase (LDH) release assay and an MTT assay–based cell viability assay to assess multiple cell lines, including HUVEC-CS, H9c2, A7r5, and C2C12 cells. The LDH release assay revealed that treatment with TP1 up to 100 µM produced no significant cytotoxicity or cell membrane-damaging effects on HUVEC-CS cells (**Fig. S2A**). In agreement with the LDH assay results, the MTT-based cell viability assay also indicated that HUVEC-CS cell viability was not altered following TP1 treatment up to 100 µM (**Fig. S2B–D**). The MTT-based cell viability assay also confirmed the nontoxic effect of TP1 at the effective dosage of 20 µM in multiple cell lines used in subsequent assays (**Fig. S2D**). In addition, the MTT-based assay indicated that TP1 has no significant effect on cell proliferation, which was validated via a flowcytometry–based cell cycle assessment (**Fig. S2C**). In agreement with our previous *in vivo* toxicity study, which did not show acute toxicity effects, e.g., cytotoxic, hemolytic, and immunogenic effects (19), the present results reflect the comparable nontoxic effect of TP1. Thus, we conclude that TP1 does not exert membranolytic, cytotoxic, or mitogenic effects at a dosage of up to 100 µM.

### TP1 alters hypoxia-modulated endothelial proteome

To study hypoxia-induced pathophysiology in ECs, we used a pSILAC proteomic method to pulse/chase the incorporation of stable isotope-labeled lysine and arginine into newly synthesized proteins (27). Briefly, HUVEC-CS cells were cultured for 24 h in SILAC medium containing ^13^C_6_ L-arginine and ^13^C_6_ and ^15^N_2_ L-lysine in the presence or absence of 20 µM TP1 [or the phosphate-buffered saline (PBS)–only control] under normal oxygenation or hypoxic conditions (**Fig. S3A, B**). After a 24 h-incubation, the cells were lysed, and their protein content was extracted for LC-MS/MS proteomic analysis, with which 7614 proteins were identified with high confidence (3 experimental replicates and a false discovery rate of <1%), including 7101 proteins identified based on more than one component peptide (Supplementary dataset). According to Pearson correlation analysis, a linear relationship (R^2^ > 0.9) existed between replicates (**Fig. S4**), indicating the high reproducibility and reliability of the experiments. All statistical analyses were conducted using data from triplicate experiments, P-values were calculated using Student’s t-test, and volcano plots were used to determine data distribution and cutoff levels (**Fig. S5A– C**). In total, 2802 proteins were significantly enriched under hypoxia, either in the presence or absence of TP1, and the relative abundance of the newly synthesized proteins was subjected to hierarchical clustering (**Fig. 2A** **and S6**). Cutoffs for differential protein synthesis were determined using log_2_ ratio distributions: ratio values of <0.76 [log2 (Ratio) < −0.4] and >1.32 [log2 (Ratio) > 0.4] for reduced and induced proteins, respectively (**Fig. S5A**). Using these cutoff values, we found that 148 and 431 proteins were induced and suppressed, respectively, under hypoxic conditions compared to normoxic conditions. We also found that TP1 treatment induced and repressed a further 39 and 172 distinct proteins, respectively, during hypoxia.

**Fig. 2:**
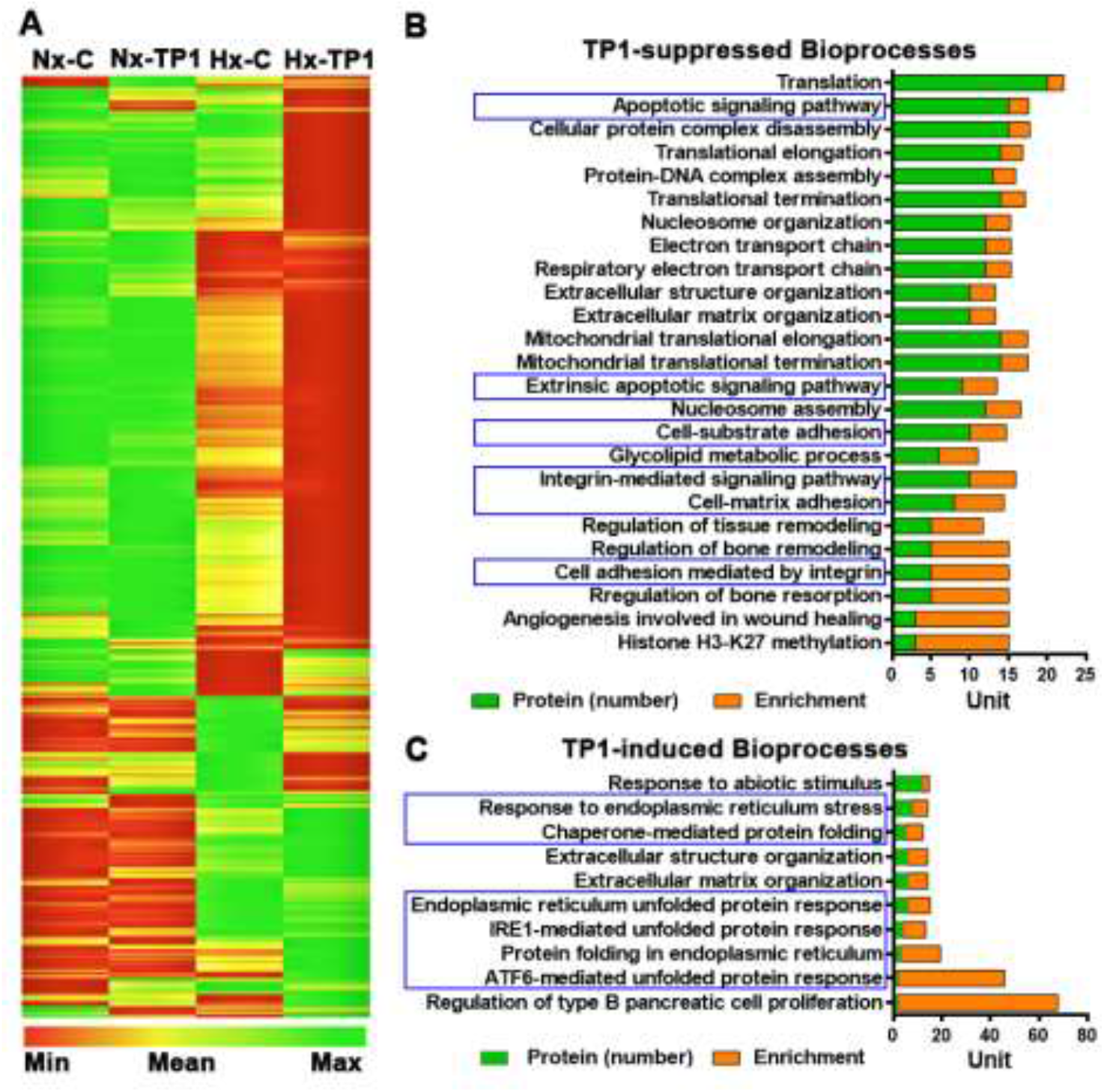
TP1-mediated differential protein synthesis in HUVEC-CS cells under normoxic and hypoxic conditions. A) Heat map representing the relative abundance of newly synthesized proteins under the indicated experimental conditions. HUVEC-CS cells were treated with 20 µM TP1 for 24 h under normoxic or hypoxic conditions, respectively, and PBS was used as the vehicle control. A SILAC–based protein labeling approach was used to label the newly synthesized proteins, which were later quantified using a LC-MS/MS–based quantitative proteomic technique. Cluster analysis of the identified proteins was performed using the online bioinformatics tool Gene Pattern (http://genepattern.broadinstitute.org) with hierarchical clustering and Pearson correlation options. Vehicle control under normoxic (Nx-C) or hypoxic (Hx-C) conditions and TP1 treatment under normoxic (Nx-TP1) or hypoxic (Hx-TP1) conditions are shown. B, C) Bioinformatics analysis of TP1-regulated bioprocesses in hypoxic HUVEC-CS cells. Figures represent TP1-suppressed bioprocesses (B) and TP1-induced bioprocesses (C) during 24 h of hypoxic stress.

### TP1 limits hypoxia-induced endothelial activation and THP-1 cell adhesion

We performed functional analyses of the proteins modulated by low-oxygen stress with or without TP1 treatment. Data analysis performed using the online bioinformatics tool GOrilla (http://cbl-gorilla.cs.technion.ac.il; 28) revealed that hypoxia induced the expression of exocytosis regulators, key mediators of leukocyte activation/inflammatory responses and various metabolic intermediaries (**Fig. S7A–C**). Importantly, TP1-treatment significantly reduced the synthesis of proteins involved in VED pathogenesis, including integrin-mediated cell adhesion molecules, cell-matrix/substrate adhesion regulators, and apoptotic signaling pathways (**Fig. 2B**). Also, hypoxia stimulates endothelial activation and promotes inflammatory responses, which are significantly reduced by treatment with TP1. Indeed, TP1 treatment also induced the synthesis of proteins associated with cellular responses to ER stress, including the ATF6-mediated unfolded protein response (UPR), IRE1-mediated UPR, and chaperone-mediated protein folding (**Fig. 2CB**). Taken together, TP1 decreases hypoxia-induced ER stress in the human endothelium.

Bioinformatics analysis of our pSILAC data indicated that hypoxia stress–activated inflammatory response pathways were active in ECs (**Fig. S7A**–**C**), leading to leukocyte activation and recruitment. Therefore, we used quantitative PCR to determine the expression patterns of leukocyte adhesion receptors during hypoxia. Our finding showed that the expression level of *ICAM1* mRNA increased 9-fold and the expression of *ALCAM*, *L1CAM*, and *VCAM1* mRNAs was also substantially upregulated (**Fig. 3D**). Analysis of proteomic data showed that ICAM1 protein synthesis was upregulated in hypoxic ECs relative to normoxic cells (**Fig. 3C**). In addition, cell adhesion assays using THP-1 human monocytic cells showed that THP-1 binding to hypoxic HUVEC-CS cells increased 2-fold (**Fig. 3A, B**). Proteomic data analysis also revealed that TP1 treatment significantly decreased the expression of ICAM1 and ALCAM at the mRNA and protein levels in hypoxic ECs (**Fig. 3C, D**) and reduced the transcription of L1CAM and VCAM1 (**Fig. 3D**). Integrins are key signaling molecules involved in the regulation of various bioprocesses such as the activation of inflammatory responses (29). The activation of integrin subsets, including αvβ3 and α5β1, is considered to trigger inflammatory responses through the activation of proinflammatory gene expression (30–32). In the present study, we found that integrin-subtype β3 synthesis was induced by hypoxia (**Fig. S8A**), which also significantly upregulated total integrin levels, including the levels of integrin subtypes α_v_, β_1_, and β_3_ in ECs (**Fig. S8B, C**). Proteomic data analysis revealed that TP1-treatment suppressed the synthesis of critical integrin subtypes in hypoxic ECs (**Fig. S8B, C**) and significantly downregulated the total expression levels of integrin subtypes α_v_, β_1_ and β_3_ in hypoxic ECs (**Fig. S8B, C**). Therefore, the TP1-mediated suppression of integrin expression might prevent the activation of integrin subtypes, including αvβ3 and α5β1, which could result in the deactivation of proinflammatory gene expression (30–32), including the expression of *ALCAM*, *ICAM1*, *L1CAM*, and *VCAM1*, in hypoxic ECs. Accordingly, THP1 cell adhesion to hypoxic ECs was decreased 2.9-fold following TP1 treatment (**Fig. 3A, B**), suggesting the high potential of TP1 affecting VED pathogenesis *in vivo*. In contrast, treatment with two major bioactive ginseng metabolites, ginsenosides Rb1 and Rg1, did not affect hypoxia-mediated proinflammatory gene activation (**Fig. 3D**), nor did these ginseng metabolites prevent hypoxia-induced leukocyte adhesion to ECs (**Fig. 3A, B**). Thus, unlike TP1, ginsenosides Rb1 and Rg1 do not have any protective impacts against hypoxia-induced endothelium activation.

**Fig. 3:**
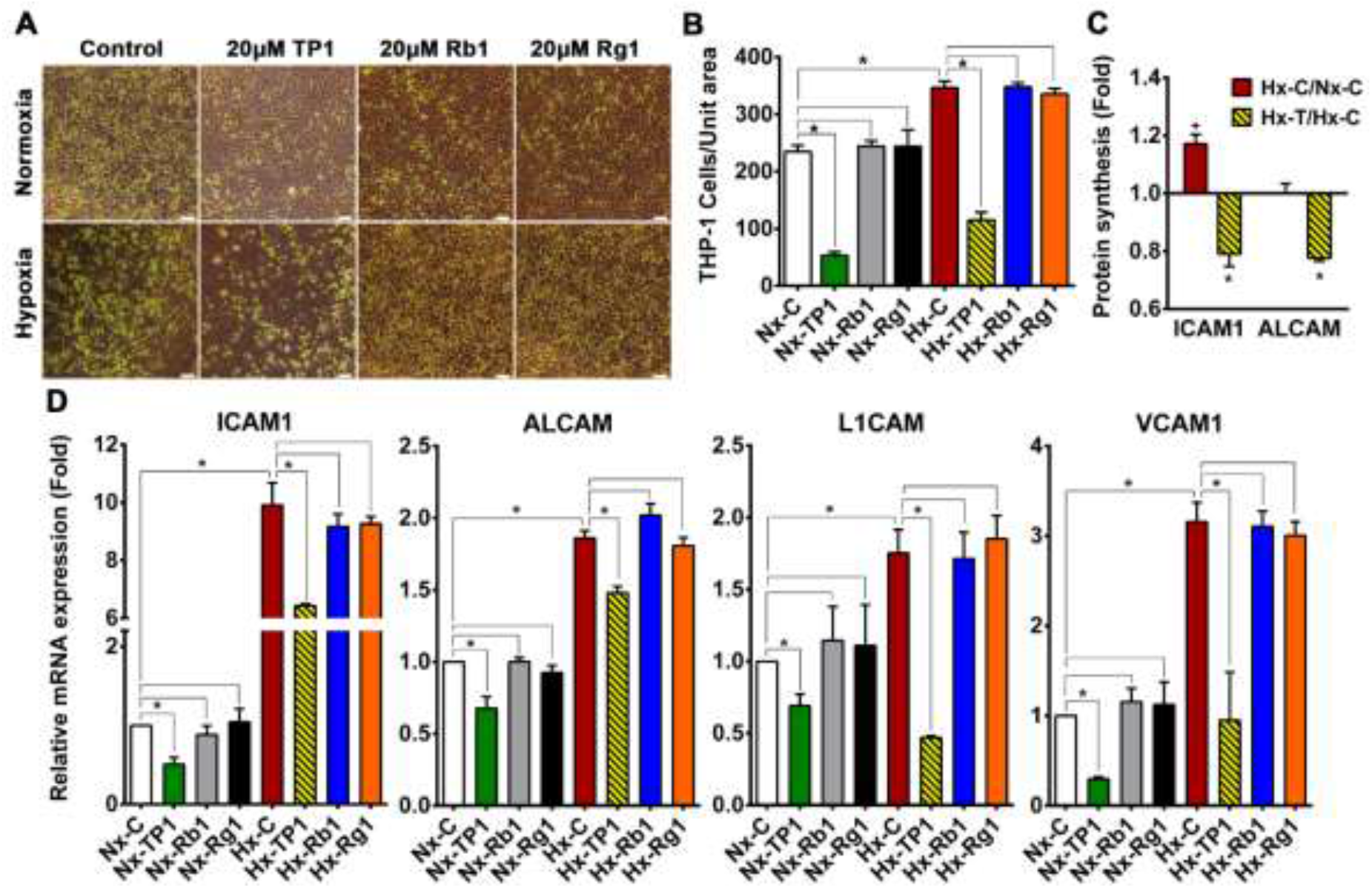
Ginsentide TP1 inhibits THP-1 cell adhesion to human vascular endothelial cells. A) Differential binding of human monocytic THP1 cells to a HUVEC-CS monolayer cultured under normoxic or hypoxic conditions in the presence of TP1/Rb1/Rg1 or the PBS vehicle–only control. Carboxyfluorescein succinimidyl ester (CFSE)-labeled THP-1 cells are shown in green. Scale bar: 100 µm. B) THP-1 adhesion was quantified via cell counting, and statistical significance was determined using triplicate experiments. C) Fold-change in the newly synthesized cell surface adhesion receptors ICAM1 and ALCAM under hypoxic conditions relative to normoxic conditions (Hx-C/Nx-C) or in TP1-treated hypoxic cells compared with vehicle-treated hypoxic cells (Hx-TP1/Hx-C). Data are means ± standard error of the mean (SEM), and statistical significance was calculated based on triplicate experiments. D) Relative mRNA expression of adhesion receptors in control and treated HUVEC-CS cells cultured under normoxic or hypoxic conditions. Data means ± SEM and statistical significance were determined using triplicate biological replicates. Vehicle control cells were cultured for 24 h under normoxic (Nx-C) or hypoxic (Hx-C) conditions and TP1/Rb1/Rg1-treated cells were cultured for 24 h under normoxic (Nx-TP1/ Nx-Rb1/ Nx-Rg1) or hypoxic (Hx-TP1/ Hx-Rb1/ Hx-Rg1) conditions, respectively. *P < 0.05.

### TP1 promotes EC survival by antagonizing apoptotic signaling during hypoxia

Given our findings that TP1 protects ECs against hypoxia-induced oxidative stress and ER stress, we assessed whether TP1 increases cell viability under hypoxic conditions. Briefly, ECs were cultured in Dulbecco’s modified Eagle medium (DMEM) containing 10% fetal bovine serum (FBS) with or without TP1 and then incubated in a normoxic or hypoxic environment for 24 h before cell survival was determined using a MTT assay. These experiments revealed that TP1 treatment increased EC viability by 20% under hypoxic conditions (**Fig. 4A, B**). Importantly, comparable protective effects were also observed in other TP1-treated cell types, including rat heart myoblast H9c2 cells (32%), rat aorta thoracic/smooth muscle A7r5 cells (31%), and mouse myoblast C2C12 cells (19%) (**Fig. 4A**). We also tested two other ginsentide members, TP3 (*P. ginseng* seed) and TP8 (*P. quinquefolius* flower), which differ from TP1 by one and three amino acid residues, respectively (**Fig. 1A**). Like TP1, TP3, and TP8 also exerted similar protective effects against hypoxic cell death in ECs (**Fig. 4B**). Collectively, these results suggest that TP1 and its closely related homologs TP3 and TP8 provide various cell types with protection against hypoxia-induced death.

Hypoxia is known to induce cell cycle arrest in the G0/G1 phase and impair mitochondrial function, leading to the apoptosis of ECs and progression to VED (33). Our proteomic data analysis showed that TP1 could suppress the synthesis of apoptosis signaling proteins in hypoxic ECs (**Fig. 4C**); therefore, we investigated the antiapoptotic effects of TP1 via the flow cytometric analysis of annexin V and propidium iodide (PI) staining. We observed a marked increase in the proportion of apoptotic and necrotic HUVEC-CS cells cultured under hypoxic conditions; however, treatment with TP1 was sufficient to abolish this increase (**Fig. 4D** **and S9A**–**C**). TP1 also conferred significant protection against cell death in an alternative doxorubicin-induced apoptosis model (**Fig. S9D, E**). This result is significant because doxorubicin is a leading drug against certain types of cancer, and TP1 exerts antiapoptotic effects to prevent doxorubicin-induced cell death during hypoxia.

We also investigated the effects of ginsenosides on the viability of ECs during hypoxic stress but found no significant effects of Rb1 or Rg1 treatment on cell survivability (**Fig. 4B**). Thus, unlike the tested ginsentides, we found these two ginsenosides have no protective effects against hypoxia-mediated cell death.

**Fig. 4.**
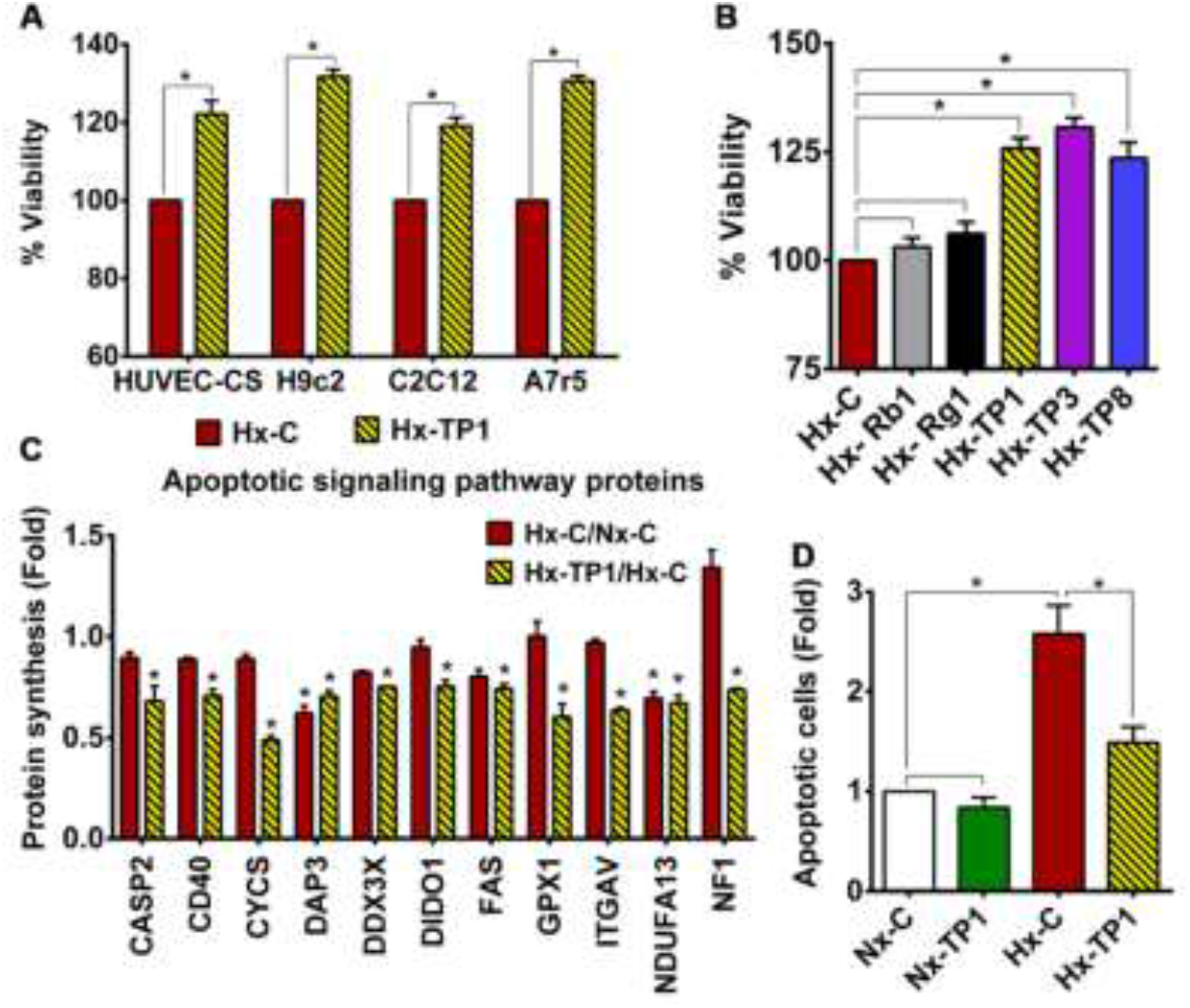
Ginsentide TP1 prevents hypoxic stress-mediated cell death. A) Viability of hypoxic cells measured using a MTT assay after treatment with TP1 or vehicle control. Relative survival is expressed as mean values, and statistical significance was calculated using three independent experimental replicates. B) Effects of ginsentides, i.e., TP1, TP3, and TP8 and ginsenosides, i.e., Rb1 and Rg1, on the viability of hypoxic HUVEC-CS cells. Viability was measured using a MTT assay after treatment with 20 µM ginsenoside (Rb1 or Rg1), 20 µM ginsentide (TP1, TP3, or TP8), or PBS (vehicle control). Relative survival is expressed as mean values ± standard error of the mean from three independent experimental replicates. *P < 0.05. C) Relative change in the expression of apoptotic signaling pathway proteins after 24 h of hypoxic culture in the presence of TP1. Statistical significance was calculated using data from triplicate experiments. C) Effect of TP1 on hypoxia-induced apoptosis in HUVEC-CS cells. Relative quantification was based on annexin V and propidium Iodide (PI) staining used to identify apoptotic cells. Statistical significance was calculated using three independent experimental replicates. Data are means ± standard error of the mean for triplicate experiments. *P < 0.05.

### TP1 treatment rescues ECs from hypoxia-induced ER stress

The disruption of ER homeostasis during hypoxia leads to the accumulation of misfolded/unfolded proteins in the ER lumen, resulting in ER stress. Hypoxia-induced ER stress initiates the unfolded protein response (UPR), which promotes cellular adaptation to stress conditions and promotes the apoptosis of cells that fail to restore normal function (34, 35). To better understand the mechanism underlying the promotion of hypoxic cell survival by TP1, we measured the mRNA expression levels of the UPR marker XBP1. We found that the spliced isoform was upregulated 20-fold in hypoxic HUVEC-CS cells compared to normoxic HUVEC-CS cells (**Fig. S10**). Under these conditions, according to our proteomic data analysis, TP1 induced the synthesis of several ER stress sensor (ATF6/IRE1)–regulated downstream proadoptive UPR genes, including those encoding chaperone proteins and autophagy proteins, in hypoxic ECs (**Fig. 5A, B**). Consistent with these proteomic findings, we also detected the upregulation of spliced-XPB1 mRNA in TP1-treated hypoxic ECs (**Fig. S10**) as well as the enhanced expression of ATF6 transcripts and multiple downstream UPR genes, e.g., *BiP* (*HSPA5*), *GRP94* (*HSP90B1*), and *EDEM* (**Fig. 5C**). Our quantitative reverse transcription–polymerase chain reaction (qRT-PCR) analyses also indicated that TP1-treatment significantly downregulated the expression of multiple genes in the PERK–eIF2a–ATF4 apoptosis pathway, including *ATF4*, *CHOP*, *DR5*, and *TRB3*, in hypoxic cells (**Fig. 5D**). Conversely, this qRT-PCR analysis showed that the reference compounds, ginsenosides Rb1 and Rg1, had no significant effect on the mRNA expression of ER stress–related genes (**Fig. S11A, B**).

Autophagy is an essential cell process, a principal component of the integrated stress response (36), and part of the ER stress-mediated proadoptive UPR response (34, 35). Indeed, western blot analysis revealed that the ratio of LC3B-II/LC3B-I increased significantly upon TP1 treatment under hypoxic conditions (**Fig. 5E, F**), indicating an increase in autophagosome formation (36). TP1-treatment also increased P62 degradation in hypoxic ECs and resulted in a lower level of P62 in these cells (**Fig. 5E, F**), further confirming the activation of autophagy. Overall, our western blot results indicated that TP1-induced selective autophagy occurred in ginsentide-treated hypoxic cells. Thus, TP1 promotes the clearance of accumulated proteins and suppresses apoptotic signaling to prevent the ER stress-induced death of hypoxic cells.

**Fig. 5.**
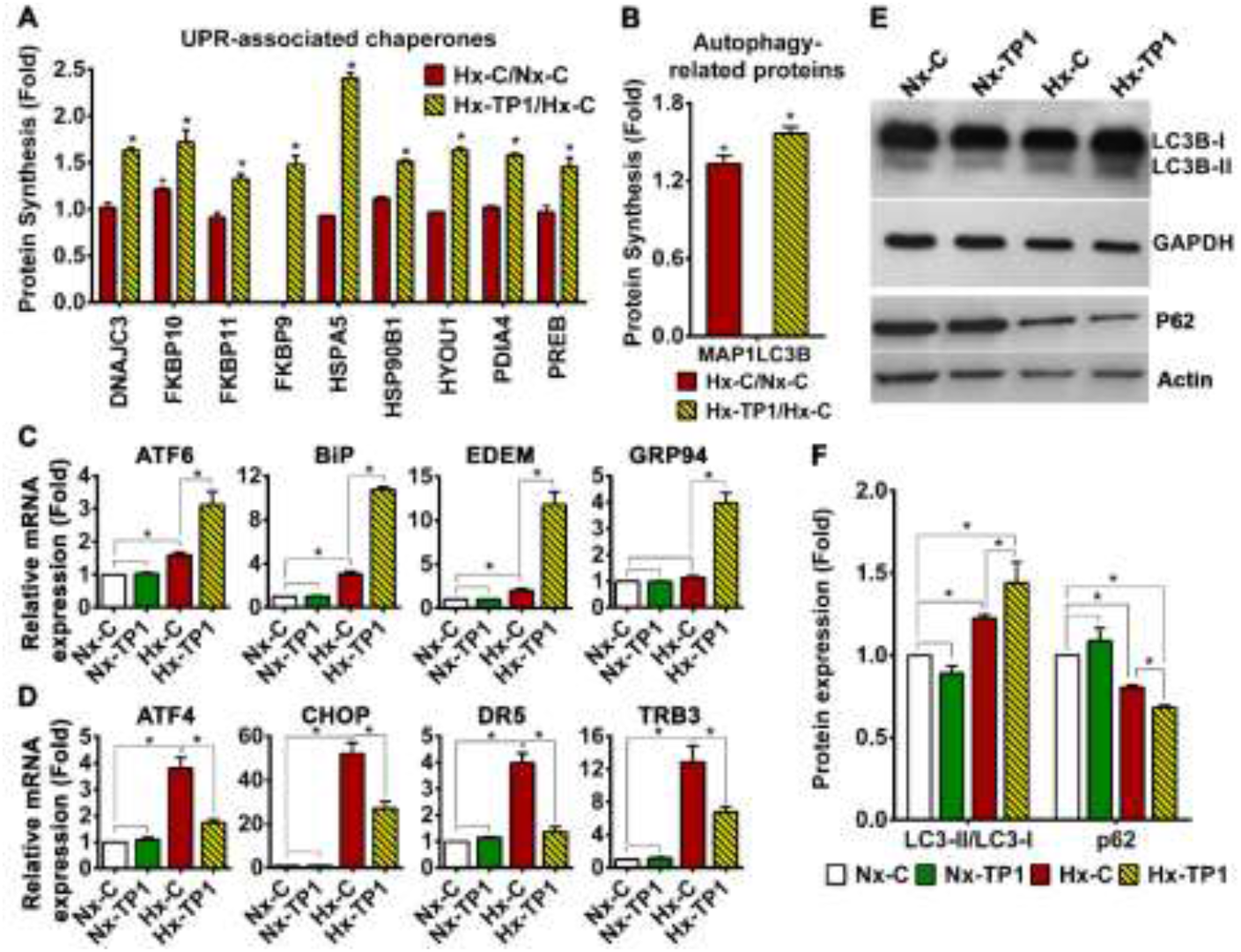
Ginsentide TP1 rescues endothelial cells from hypoxia-induced ER stress. A-B) Relative expression levels of UPR-associated chaperone proteins (A) and autophagy-related proteins (B) under hypoxic conditions compared with normoxic conditions (Hx-C/Nx-C) or in TP1-treated hypoxic cells relative to PBS-treated hypoxic cells (Hx-TP1/Hx-C). Data are means ± SEM, and statistical significance was calculated using triplicate experiments. Significance *P < 0.05 respect to Nx-C and ^+^P < 0.05 respect to Hx-C. C, D) Relative mRNA expression of ER stress–related genes in HUVEC-CS cells cultured under various conditions. Data are means ± SEM, and statistical significance was calculated using triplicate biological replicates. D, E) Western blot images of autophagy markers, including LC3B-I and II and P62 (E), under the indicated experimental conditions. F) Relative quantification of LC3B-II/LC3B-I ratio and relative expression of P62 under the given experimental conditions (quantified based on western blot band intensity). Data are means ± SEM, and statistical significance was calculated using triplicate biological replicates. Vehicle control cells were cultured for 24 h under normoxic (Nx-C) or hypoxic (Hx-C) conditions and TP1-treated cells were cultured for 24 h under normoxic (Nx-TP1) or hypoxic (Hx-TP1) conditions. *P < 0.05.

Indeed, when we measured protein aggregation using thioflavin-T staining and fluorescence microscopic analysis, we found a 1.6-fold increase in protein accumulation in hypoxic HUVEC-CS cells, which was effectively reversed by TP1 treatment (**Fig. 6A, B**). The protective effect of TP1 against hypoxia was also confirmed using alternative cell lines, i.e., H9c2, A7r5, and C2C12, in which the reduction in thioflavin-T staining upon TP1-treatment was comparable to that in HUVEC-CS cells (**Fig. 7A, B**). Consistent with the fluorescence-based analyses, the ultracentrifugation and quantitation of protein aggregates confirmed that protein accumulation in HUVEC-CS cells during hypoxic stress was markedly reduced by TP1 treatment, suggesting that TP1 is an efficient stimulator of the UPR (**Fig. 6C**).

**Fig. 6.**
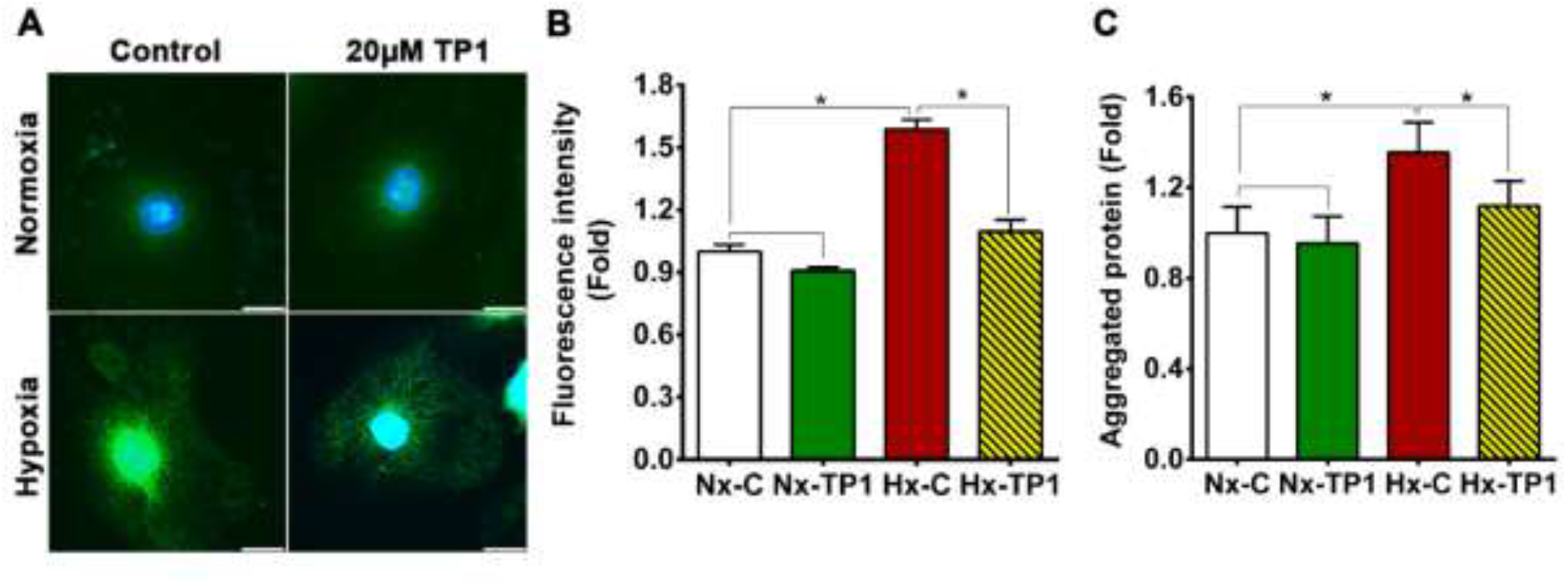
TP1-mediated recovery of ER/protein homeostasis in hypoxic endothelial cells. A) Thioflavin-T (ThT) staining of protein aggregates in HUVEC-CS cells cultured under the indicated conditions. Aggregated proteins were stained with ThT (green), and nuclei were counterstained with DAPI (blue). Scale bar: 20 µm. B) Relative quantification of protein aggregation in cells subjected to the indicated experimental conditions. Protein aggregation was quantified by calculating the average fluorescence intensity of about100 individual cells from four experimental replicates. C) Relative abundance of aggregated proteins isolated from HUVEC-CS cells cultured under different conditions. Data are means ± SEM of independent experimental triplicates. Vehicle control cells were cultured for 24 h under normoxic (Nx-C) or hypoxic (Hx-C) conditions, and TP1-treated cells were cultured for 24 h under normoxic (Nx-TP1) or hypoxic (Hx-TP1) conditions. *P < 0.05.

**Fig. 7.**
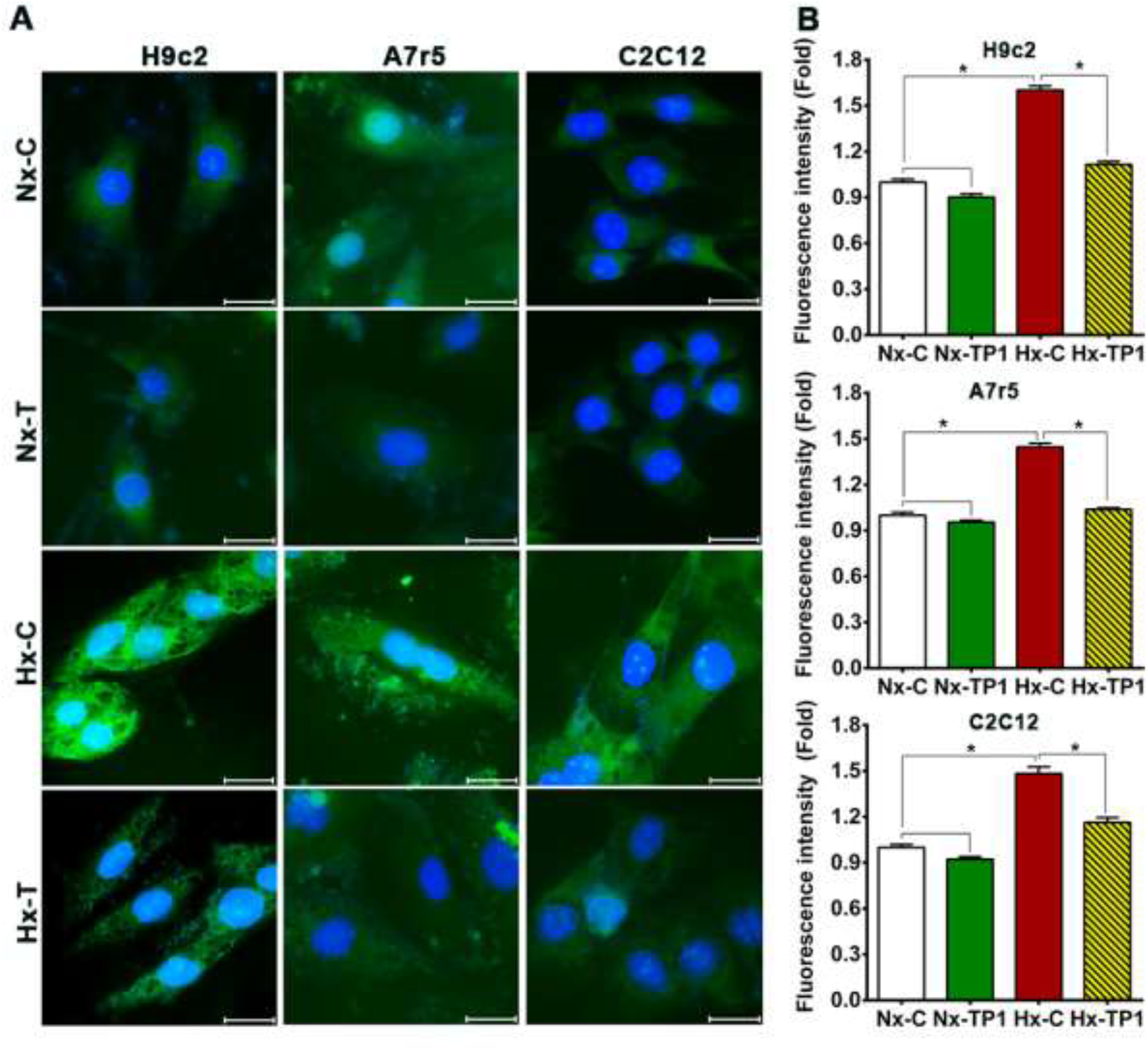
TP1-mediated recovery of ER/protein homeostasis in hypoxic cardiovascular cell types. A) Thioflavin-T (ThT) staining of aggregated proteins in the cardiovascular cell lines H9c2, A7r5, and C2C12 cultured under various experimental conditions. Protein accumulation was visualized by staining with ThT (green), and nuclei were counterstained with DAPI (blue). Scale bar: 20 µm. B) Relative quantification of protein accumulation under different experimental conditions was performed by calculating the average fluorescence intensity of >100 individual cells from four experimental replicates. Vehicle control cells were cultured for 24 h under normoxic (Nx-C) or hypoxic (Hx-C) conditions and TP1-treated cells were cultured for 24 h under normoxic (Nx-TP1) or hypoxic (Hx-TP1) conditions. *P < 0.05.

### TP1 restores NO production and reduces reactive oxygen species generation in hypoxic ECs

Under normal physiological conditions, NO maintains the quiescent state of the endothelium, while disruption of NO signaling can trigger the endothelial expression of mediators that promote leukocyte adhesion and inflammation (37). Under oxidative stress conditions, the reaction of NO with superoxide anions generates peroxynitrite, which significantly impairs NO production by nitric oxide synthases, thereby promoting endothelial activation and progression to VED (37–39). Given our finding that TP1 limits endothelial activation under hypoxic conditions, we assessed whether TP1 acts via effects on the generation of NO and reactive oxygen species (ROS). Using a fluorogenic staining approach, i.e., a 2′,7′-dichlorofluorescein diacetate (DCFDA) assay, we found that intracellular ROS levels were higher in hypoxic ECs than normoxic cells; however, TP1-treatment reduced ROS levels by 1.4-fold in hypoxic ECs (**Fig. 8AB**). Similarly, a 1.3-fold reduction in intracellular NO was detected in hypoxic ECs, but this deficit was effectively reversed by TP1 treatment (**Fig. 4B**). Accordingly, when the expression of eNOS was assessed using western blotting, protein levels that were reduced under hypoxic conditions were partially restored by TP1 treatment (**Fig. 8C**). Consistent with these data, hypoxic stress was found to induce a 1.3-fold decrease in eNOS phosphorylation at residue Ser^1177^, which is known to modify enzyme activity under both physiological and pathological conditions (40, 41); however, TP1-treatment reversed this deficiency in eNOS phosphorylation (**Fig. 8C**). Taken together, our data indicate that TP1 limits ROS generation and maintains NO bioavailability to prevent endothelial activation under hypoxic conditions. Bioactive ginsenosides are known to counteract redox regulation, promote NO production, and prevent ROS generation; thus, they protect cells against oxidative damage (25, 42–44). Consistent with these reported effects, we found that treatment with the ginsenosides Rb1 and Rg1 reduced the intracellular ROS levels of both normoxic and hypoxic ECs, comparable with the impact of TP1-treatment (**Fig. S12A, B**).

**Fig. 8:**
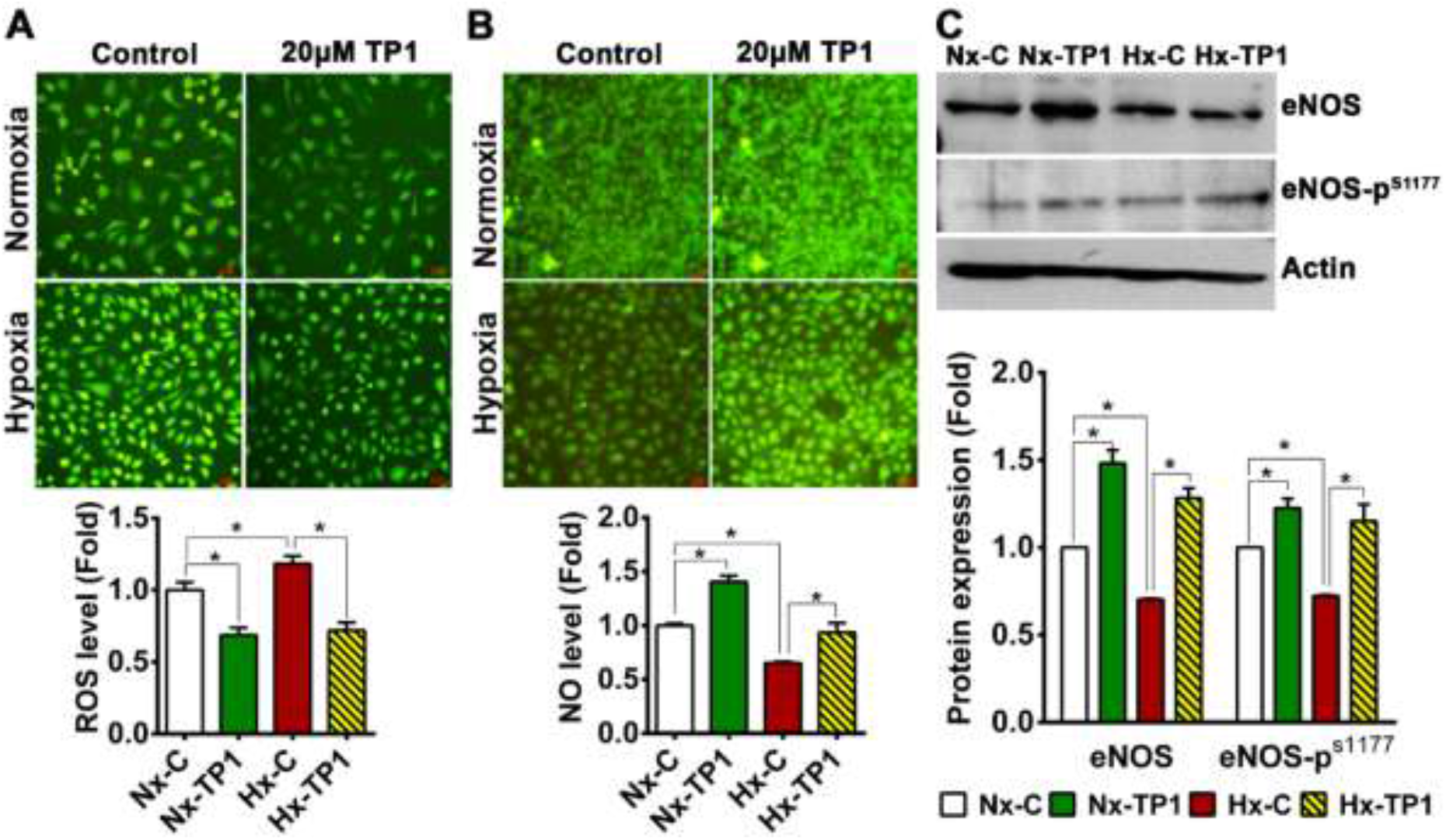
Ginsentide TP1 reduces intracellular reactive oxygen species (ROS) generation and restores nitric oxide (NO) generation in endothelial cells. A) Effect of TP1 on ROS levels in HUVEC-CS cells cultured under different experimental conditions. ROS activity was measured by (DCFDA)-based fluorogenic assay. Scale bar: 25 µm. Relative quantification was performed using the relative fluorescence intensity. Statistical significance was calculated using five biological replicates. B) HUVEC-CS cells were cultured with or without TP1/Rb1 /Rg1 for 24 h under normoxic or hypoxic conditions, and NO levels were measured via DAF-2DA staining. Scale bar: 25 µm. Relative quantification of NO was based on the fluorescence intensity in each image, and statistical significance was calculated using four biological replicates. C) Relative expression of eNOS and Ser^1177^-phosphorylated eNOS (eNOS-p^S1177^) under the indicated experimental conditions (quantified based on western blot band intensity). Data are shown as means ± standard error of the mean, and statistical significance was calculated using biological triplicates. Vehicle control cells were cultured for 24 h under normoxic (Nx-C) or hypoxic (Hx-C) conditions, and TP1/Rb1 /Rg1-treated cells were cultured for 24 h under normoxic (Nx-TP1/ Nx-Rb1/ Nx-Rg1) or hypoxic (Hx-TP1/ Hx-Rb1/ Hx-Rg1) conditions. *P < 0.05.

## Discussion

Ginseng, the most valuable herb in traditional Chinese medicine, is prescribed to treat a wide range of disorders, including cardiovascular disease (17, 18, 45). Thus far, ginsenosides, which belong to the small-molecule metabolites of the triterpene saponin family, have received the most attention as the active compounds to account for the ethnomedicinal effects of ginseng (45, 46). Despite an abundance of ginseng research, ginsenosides alone cannot adequately explain the ‘cure-all’ effect of ginseng in its ethnomedicinal uses (45, 46). Recently, we reported that TP1 coordinates multiple systems to exert anti-stress and cardiovascular benefits (25). This report shows that ginsentide TP1 plays an important adaptogenic role in maintaining protein homeostasis in hypoxic cells and suppressing ER-stress-induced apoptosis.

Ginsentides have two distinct advantages over most unconstrained peptide biologics of comparable size. First, ginsentides, containing about 25% cysteine, are highly cystine-braced, rendering them structurally compact and highly resistant to proteolytic degradation. Second, ginsentide TP1, the representative and most abundant ginsentide is a cell-penetrating peptide and can target intracellular protein-protein interactions. Also, TP1 does not exhibit any toxic or membrane-damaging effects. In comparison, ginsenosides have poor bioavailability, thereby limiting their clinical applications (47). TP1. The cell-penetrating property can also be found in certain highly cystine-braced microproteins, such as the plant-derived 8C-hevein-like peptides (20–24, 26). Thus, hyperstability and cell-penetrating properties of ginsentides are desirable attributes for drug development.

The vascular endothelium is comprised of a monolayer of cells that maintain hemostasis by regulating vascular tone, cell adhesion, smooth muscle proliferation, and tissue inflammation. The dysregulation of these mechanisms leads to VED, which can result in systemic inflammatory pathology, adverse thrombotic events, and vascular tissue remodeling. A link between hypoxia and VED has been established, as well as their consequences in relation to cardiovascular disorders (1–6). Because preventing hypoxia-linked VED will be beneficial for treating cardiovascular disorders, we established a pulsed SILAC–based quantitative proteomic approach to determine the effects of TP1 on hypoxia-linked VED. We profiled the newly synthesized proteome of normoxic and hypoxic endothelium cells with or without TP1 treatment and analyzed the differentially expressed newly synthesized proteins. Our findings showed that hypoxia and TP1 treatment alters the expression profile of newly synthesized proteins in endothelium cells. Gene ontology analysis revealed that hypoxia promotes the expression of endothelium dysfunction–linked pathway proteins. In addition, TP1-treatment reduced the expression of VED pathogenesis-linked pathway proteins, including integrins, cell adhesion molecules, and apoptotic signaling proteins. In this regard, we showed that pulsed SILAC–based quantitative proteomics is useful in the functional characterization of cell-penetrating and cystine-dense microproteins such as ginsentides.

Hypoxia promotes inflammatory signaling (9) in ECs, resulting in the increased surface expression of adhesion molecules, including ICAM1 and VCAM1, which promote leukocyte/platelet binding and progression to VED (48, 49). Integrin complexes, including αvβ3, are involved in the regulation of surface adhesion molecules (30–32) and the TP1-mediated suppression of integrin αv, β1, and β3 expression, which lead to decreased surface molecule expression levels in hypoxic ECs and reduced-THP-1 cell adhesion to these cells. In the present study, hypoxic ECs enhanced the expression of adhesion molecules and increased leukocyte binding by 2-fold. In contrast, hypoxic ECs treated with TP1 exhibited reduced adhesion molecule expression at both the mRNA and protein levels. Accordingly, leukocyte binding to hypoxic ECs was decreased by 2.9-fold in the presence of TP1, indicating the potent effects of ginsentides on the activation of the endothelium (50, 51). In contrast, ginsenoside Rb1 and Rg1, two bioactive components of ginseng, displayed no notable effects on the surface molecule expression and THP-1 cell adhesion of ECs.

Cellular stress, including hypoxia, can cause defects in protein folding that leads to protein aggregation in the ER lumen and induce ER stress (52, 53), which is a crucial component of endothelial dysfunction and cardiovascular disease (15, 54, 55). The eNOS/NO pathway plays a critical role in maintaining the intracellular redox balance and protecting against ER stress in ECs (56). Hence, the disruption of this pathway results in oxidative stress and proinflammatory responses that promote VED (15, 57). One mechanism underlying the protection of hypoxic cells against ER stress is the evolutionarily conserved UPR, which first attempts to restore protein homeostasis and later triggers apoptosis if recovery is unsuccessful (34, 35, 58). The UPR plays a vital role in determining the fate of stressed cells. In the current study, hypoxic HUVEC-CS cells exhibited the 18-fold upregulation of spliced XBP1 mRNA, an archetypal marker of UPR activation (59). Indeed, low-oxygen conditions also promoted the accumulation of unfolded proteins in multiple cell types, including ECs, myocytes, cardiomyocytes, and vascular smooth muscle cells, and the significant upregulation of the cell death signaling genes *ATF4*, *CHOP*, *DR5*, and *TRB3*. However, treatment with TP1 increased the hypoxic EC expression of anti-aggregation ERED genes and molecular chaperones that support protein folding, including *DNAJC3*, *HSP90B1*, *HSPA5*, and *HYOU1*, which help restore protein homeostasis and reduce ER stress in hypoxic ECs. In addition, TP1 promoted selective autophagy as a part of the proadaptive UPR response (35, 36), thereby enhancing protein clearance and restoring protein homeostasis in hypoxic ECs. Our proteomic results revealed that the expression of apoptosis signaling proteins and cell death rates were significantly reduced following TP1 treatment in hypoxic ECs, myocytes, cardiomyocytes, and vascular smooth muscle cells. Taken together, our data indicate that TP1 exerts potent effects on the UPR pathway and reduces ER stress-induced apoptotic signaling, thereby increasing overall EC survival under hypoxic conditions (**Fig. 9**). The ability of TP1 to protect ECs from stress-induced apoptosis could be exploited clinically to treat various human disorders, particularly hypoxia-associated cardiovascular disorders.

Hypoxia has been shown to impair the activity of the NO biosynthetic enzyme eNOS via multiple mechanisms, including the subduction of protein phosphorylation at Ser^1177^ (40, 41, 60, 61), which leads to the disruption of NO-mediated protective pathways and endothelial activation by redox signaling (60, 62). Loss of eNOS function reduces NO production and increases superoxide radical generation in hypoxic ECs, leading to endothelium activation. However, in the present study, we found a significant decrease in eNOS expression and NO levels, in parallel with elevated ROS production, in hypoxic ECs. Importantly, TP1 treatment reversed this hypoxia-induced effect in hypoxic ECs, increasing eNOS expression and phosphorylation at Ser^1177^, which led to the recovery of NO synthesis. The TP1-mediated restoration of the redox balance in hypoxic cells via the eNOS/NO pathway may prevent endothelium dysfunction (25, 50, 51). Notably, the tested ginsenosides Rb1 and Rg1 exerted a comparable effect (25), i.e., they restored the redox balance by increasing the NO level and reducing the ROS load in hypoxic ECs.

**Fig. 9.**
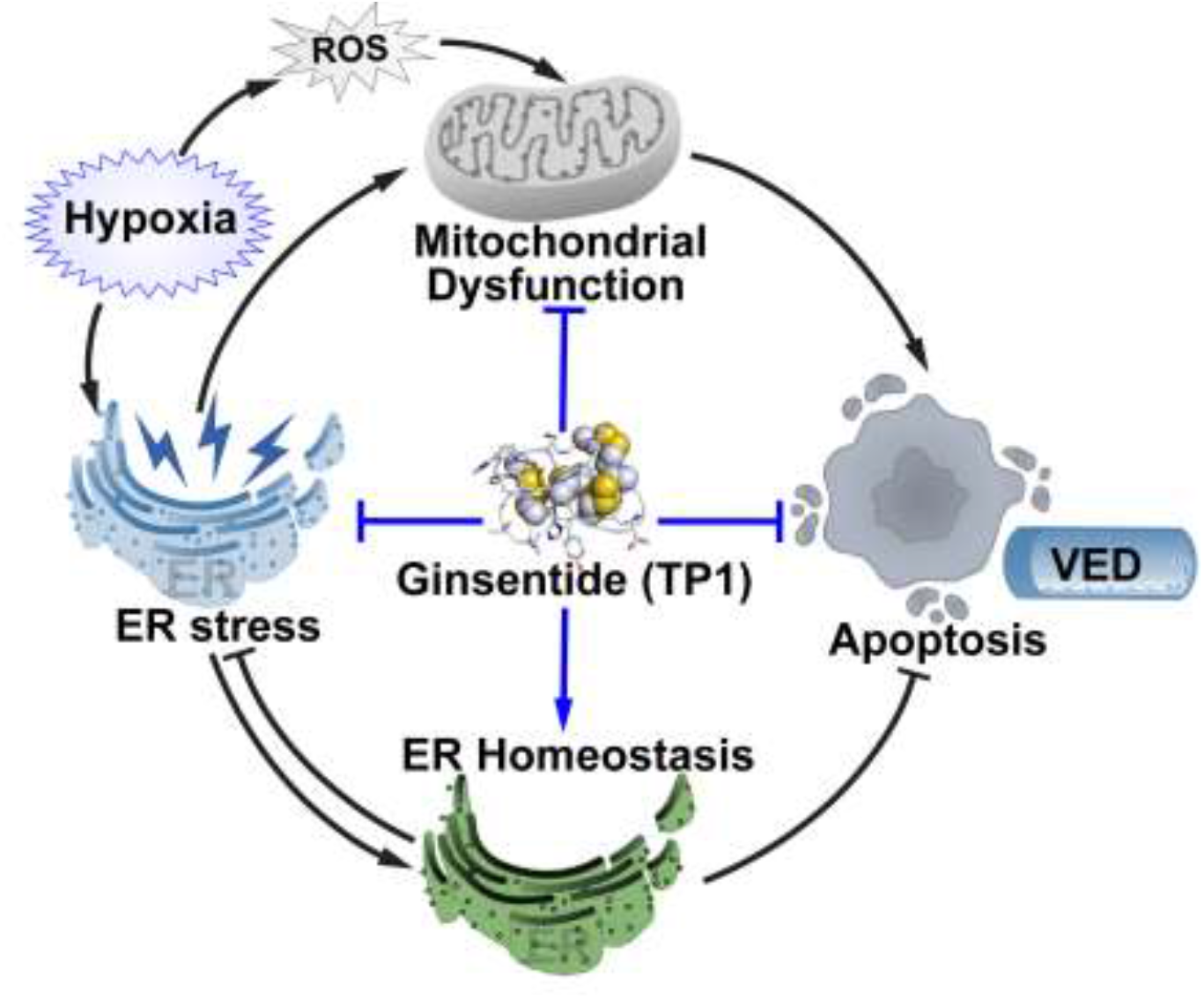
Proposed model for TP1-mediated protective effects against hypoxic stress in the endothelium. Endoplasmic reticulum (ER) stress initiates the unfolded protein response (UPR) as a part of the integrated stress response. UPR promotes both cellular adaptations to stress conditions and the apoptosis of cells that fail to restore normal function. Adaptive UPR prevents ER stress by restoring ER/ protein homeostasis. In contrast, prolonged ER stress induces mitochondrial dysfunction and promotes apoptotic cell death. Hypoxia-induced ER stress in endothelium cells initiates proadaptive responses. Prolonged hypoxia leads to prolonged ER stress, which is proposed to cause the failure of proadaptive responses and ensure apoptotic cell death, leading to endothelium dysfunction. Ginsentide TP1 is thought to minimize the effect of ER stress in hypoxic endothelial cells by restoring ER/protein homeostasis via the activation of proadaptive responses and suppression of apoptotic cell death.

## Conclusion

Hypoxia impairs essential cellular functions to promote endothelial cell activation and progression to VED. Low oxygen conditions also induce ER stress, which activates apoptotic signaling in hypoxic endothelial cells and multiple other cell lineages affected by cardiovascular pathology. Ginsentide TP1 rescues protein homeostasis in hypoxic cells and suppresses ER-stress-induced apoptotic signals to improve the survival of multiple cardiovascular cell types. TP1 also restores NO bioavailability and reduces intercellular ROS in hypoxic endothelial cells via restorative effects on the NO/eNOS pathway, which also protects against hypoxia-induced VED. Our results provide strong support that ginsentides are unexplored microproteins responsible for the ethnomedicinal benefits of ginseng in cardiovascular diseases. Also, ginsentides could provide therapeutic leads useful in treating patients at risk of VED and associated cardiovascular complications.

## Experimental procedures

### Reagents

All chemicals and reagents were purchased from Sigma-Aldrich (USA) unless otherwise indicated. Antibodies against GAPDH (6C5) and actin (C-2) were purchased from Santa Cruz Biotechnology (Santa Cruz, USA), whereas anti-LC3B (L7543) antibody was purchased from Sigma-Aldrich. Anti-eNOS (D9A5L), anti-phospho-eNOS (Ser1177), anti-integrin αV (D2N5H), anti-integrin β1 (D6S1W), anti-integrin β3, HRP-linked anti-mouse IgG, and HRP-linked anti-rabbit IgG antibodies were obtained from Cell Signaling Technology (USA). Clarity Max Western ECL Blotting Substrate (1705062) was purchased from Bio-Rad Laboratories (Italy).

### Extraction and purification of TP1 and other ginsentides

Peptide extraction was performed using the method described by Tam *et al.* (19). Briefly, dried *P. ginseng* flowers were pulverized and blended with Milli-Q water (100 g/L). The extract was then centrifuged, and the obtained supernatant was filtered through a 0.22 μm membrane (Thermo Fisher Scientific, MA, USA). Purification was performed using multiple rounds of reversed-phase high-performance liquid chromatography (RP-HPLC), which was conducted using a C18 column (250 × 22 mm; 5 μm particle size; Phenomenex, USA) at a flow rate of 5 mL/min with a linear gradient of 1%/min of 10%– 80% buffer B [0.1% trifluoroacetic acid (TFA) in acetonitrile (ACN)] and buffer A (0.1% TFA in HPLC water). Final purification was performed via analytical RP-HPLC using a C18 column (250 × 4.6 mm; 5 μm particle size; Phenomenex) at a flow rate of 1 mL/min using the same gradient. MALDI-TOF mass spectrometry was used to identify the peptide-containing fractions. Ginsentide TP3 and TP8 were isolated from dried *P. ginseng* seeds and dried *P. quinquefolius* flowers, respectively, and purified using the same method.

### Cell culture

HUVEC-CS cells, rat cardiomyoblast cells (H9c2), rat aorta thoracic/smooth muscle cells (A7r5), and mouse myoblast cells (C2C12) were maintained in DMEM supplemented with 10% FBS at 37°C in 5% CO2. THP1 cells were maintained in an RPMI medium supplemented with 10% FBS at 37°C in 5% CO2.

### Confocal microscopic-based cellular uptake analyses

Cy3–TP1 was used to study the cellular uptake of TP1 in HUVEC-CS cells. To visualize the intracellular distribution of Cy3–TP1, cells were grown on an 8-well chamber slide (Ibidi, Germany) overnight before they were incubated with 1 μM Cy3–TP1 at 37°C for 1 h and counterstained with LysoTracker and Hoechst 333241. Postincubation, the medium was drained, and the cells were washed gently with PBS. Fresh medium was added prior to imaging. Slides were observed and imaged using a Zeiss LSM 710 confocal microscope.

### Pulsed SILAC labeling

For pSILAC labeling, 2 × 10^6^ HUVEC-CS cells were seeded onto 10 cm Petri dishes and cultured overnight at 37°C in 5% CO_2_. The supernatant was aspirated the following day and replaced with SILAC medium containing ^13^C_6_ L-arginine and ^13^C_6_ and ^15^N_2_ L-lysine together with 10% SILAC-compatible dialyzed FBS. The cells were then supplemented with 20 µM TP1 or PBS vehicle control, after which the dishes were transferred into hypoxia chambers from which the air was removed by flashing the chambers with hypoxic gas (95% N_2_, 5% CO_2_, and <0.1% O_2_) for 10 min at a constant flow rate of 15 L/min. The chambers were then sealed and incubated at 37°C for 24 h. Control cells were instead maintained under standard culture conditions (i.e., normoxia: 21% O_2_ and 5% CO_2_) at 37°C for the 24 h incubation period. Subsequently, the cells were trypsinized and washed three times with ice-cold PBS before they were stored at −80°C until they were <u>used</u> in proteomic experiments. The entire experimental protocol was performed in triplicate, as summarized in **Fig. S2B**.

### Protein extraction and tryptic digestion

The cell pellets obtained from each experimental condition were suspended in lysis buffer (8 M urea in 100 mM ammonium bicarbonate supplemented with protease inhibitor cocktail) and sonicated at 30% amplitude for the 30 s. The cell lysates were clarified via centrifugation at 20,000 × *g* for 10 min, and the protein content of the lysates was quantified using the Bradford protein assay. For each sample, 500 μg of protein was diluted with 100 mM ammonium bicarbonate buffer to decrease the urea concentration to <1 M. The samples were then reduced using 10 mM dithiothreitol at 37°C for 2 h, after which they were alkylated with 55 mM iodoacetamide at room temperature for 45 min in the dark. Finally, the samples were digested via incubation with trypsin (with an enzyme:protein ratio of 1:50) overnight at 37°C, after which the reaction was stopped by adding formic acid to a final concentration of 0.5%. Tryptic peptides were then desalted using Sep-Pak C18 cartridges (Waters, Milford, MA, USA), vacuum dried, and stored for HPLC fractionation.

### Reverse-phase chromatography fractionation and liquid chromatography with tandem mass spectrometry

Tryptic peptides were fractionated using reverse-phase chromatography separation performed at pH 8. Briefly, the desalted tryptic peptides were reconstituted in buffer A (0.02% NH4OH in water) and fractionated using an X-Bridge C18 column (4.6 × 200 mm; 5 μm particle size; 130 Å pore size; Waters) at a flow rate of 1 mL/min in an HPLC unit (Prominence, Shimadzu, Kyoto, Japan). The gradient was as follows: 100%– 95% buffer A for 3 min; 5%–35% buffer B (80% ACN and 0.02% NH4OH) for 40 min; 35%–70% buffer B for 12 min; and 70% buffer B for 5 min. The 280 nm wavelength was selected for chromatogram recording, and 64 fractions were collected within a 65 min period. The collected fractions were combined into 13 pools using a concatenated pooling strategy, after which they were dried via vacuum centrifugation. The samples were then subjected to liquid chromatography with tandem mass spectrometry (LC–MS/MS) analysis using a Q Exactive mass spectrometer coupled with an online Dionex Ultimate 3000 RSLC nano-LC system (Thermo Fisher Scientific). Peptide samples were separated using a Dionex EASY-spray column (PepMap C18; 3 μm; 100 Å) and sprayed through an EASY nanospray source at a voltage of 1.5 kV. The range 350–1600 m/z was used for the full MS scan with a resolution of 70,000 at m/z 200. The maximum ion accumulation time was set to 100 ms, and dynamic exclusion was set to 30 s. The top 10 ions above a threshold of 1000 counts were selected for high-energy collision dissociation fragmentation using 28% normalized collision energy with a maximum ion accumulation time of 120 ms as a fragmentation parameter. The other parameters were set as follows: an AGC setting of 1e + 06 for the full MS scan and 2e + 05 for the MS2 scan, an isolation width of 2 Da for the MS2 scan, the exclusion of single and unassigned charged ions from the MS/MS scan, and an underfill ratio of 0.1%. All raw data were acquired using Xcaliber 2.2 (Thermo Scientific). Samples from triplicate experiments were run as technical duplicates.

### Mass spectrometric data analysis and bioinformatics

Protein identification and quantification were performed using MaxQuant version 1.5.2.8 (Max Planck Institute of Biochemistry). The parameters used were as follows: carbamidomethyl at cysteine as a static modification; N-term acetyl, methionine oxidation, and asparagine and glutamine deamidation as dynamic modifications; trypsin/P and a maximum of two missed cleavages as digestion parameters; and a 30-ppm precursor mass and 0.5 Da fragment mass tolerance. The UniProt Knowledgebase of human proteins (downloaded on July 25, 2016, including 70,849 sequences and 23,964,784 residues) was used as a search database with a false discovery rate cutoff of <0.01% both at the peptide and protein levels. For SILAC quantification, Arg6 and Lys8 were set as labeling residues. Searched data were extracted in text format for further analysis. Technical duplicates were searched in combination, and statistical analysis was conducted on data collected in triplicate experiments. The online bioinformatics tool GOrilla was used for bioinformatics analyses (28).

### LDH assay

A LDH release–based cytotoxicity assay was conducted using a CytoSelect LDH Cytotoxicity Assay Kit (CBA-241, Cell Biolabs, Inc., CA, USA). Briefly, HUVEC-CS cells were cultured with or without 100 µM TP1 for 24 h at 37°C in 5% CO_2_. PBS was used as a vehicle control, and 1% Triton X-100 was used as a positive control, i.e., as a cell death and membrane-damaging agent. The culture medium was collected, transferred, and mixed with the LDH assay reagent at a ratio of 9:1. The reaction mixture was then incubated at 37°C for 2 h. Quantification was performed calorimetrically using a wavelength of 450 nm. The assay was conducted in triplicate, and statistical analyses and graphing were performed using GraphPad Prism (version 6.01).

### MTT assay

Cells were cultured in DMEM with or without 20 µM TP1 (or dose maintained) under normoxic and hypoxic conditions for 24 h, after which MTT reagent (final concentration of 0.5 mg/mL) was added, and the cells were incubated for a further 2 h at 37°C. The culture medium was then aspirated, and formazan crystals were dissolved in dimethyl sulfoxide to conduct colorimetric quantification at 570 nm using a microplate reader (Tecan Magellan, Switzerland) with a reference wavelength of 630 nm. Experiments were conducted in triplicate, and the resultant data were used for statistical analysis. For ginsentide TP3 and TP8 and ginsenoside Rb1 and Rg1 treatments, cells were treated with 20 µM of the respective compounds for 24 h under normoxic and hypoxic conditions, respectively, with PBS used as a vehicle control. Cell viability was assessed using a MTT assay, as described.

### Monocyte–epithelium adhesion assay

HUVEC-CS cells were grown as a monolayer in DMEM supplemented with 20 µM TP1 (or the PBS control) under normoxic or hypoxic conditions for 24 h. The ginsenosides Rb1 and Rg1 were used as reference controls at 20 µM. THP-1 cells were cultured in RPMI medium supplemented with 10% FBS. Immediately prior to their use in adhesion assays, the THP-1 cells were harvested, washed three times with PBS, and labeled with 10 µM carboxyfluorescein succinimidyl ester (CFSE) for 10 min at 37°C. The cell culture medium was aspirated from the HUVEC-CS monolayers, which were then washed with a serum-free RPMI medium. CFSE-labeled THP1 cells were then seeded onto the HUVEC-CS monolayer at a density of 2 × 10^5^ cells/well in serum-free RPMI medium and incubated for 90 min at 37°C. After incubation, the culture medium was drained, and the HUVEC-CS monolayer was washed three times with PBS to remove nonadherent leukocytes. Images of adherent leukocytes were then captured using the green fluorescence channel of a Nikon ECLIPSE Ti-S inverted microscope system (Nikon, Kanagawa, Japan). Cells were quantified via cell counting, and statistical analyses were performed using data collected in triplicate experiments.

### NO detection

Intercellular NO levels were measured using a diaminofluorescein-2 diacetate (DAF-2DA)–based assay. ECs were cultured under various experimental conditions for 24 h before the assay began. For hypoxic cultures, the cells were also treated with 100 µM CoCl_2_. First, the culture medium was aspirated, after which the cells were washed with Hank’s Balanced Salt Solution. The cells were incubated with 1 μM DAF-2DA for 60 min at 37°C, after which the medium was refreshed, and images of labeled cells were captured using the green fluorescence channel of the Nikon ECLIPSE Ti-S inverted microscope system. Fluorescence intensity was quantified by analyzing the labeled cells via ImageJ. The experiment was performed in triplicate.

### Western blot analysis

First, 35 µg of each protein sample was resolved on a 10% acrylamide gel and transferred onto a PVDF membrane. Immunoblotting was then performed using antiprotein antibodies, and detection was achieved using an ECL system (Millipore, MA, USA).

### Cell cycle analysis

Cells cultured under different experimental conditions were harvested via trypsinization and washed with ice-cold multivalent cation-free PBS. The cells were then fixed in ethanol at −20°C for 24 h and stained with 0.5 mg/mL propidium iodide for 15 min at 37°C. DNA content was measured using a BD FACSCalibur flow cytometer (BD Biosciences, CA, USA), and data analysis was performed using Weasel (version 3.3.2).

### qRT-PCR

RNA was extracted from cells using RNA isolation kits (Macherey-Nagel, Duren, Germany) according to the ’manufacturer’s protocol. RevertAid H Minus First Strand cDNA Synthesis Kits (Thermo Fisher Scientific) were used to synthesize first strand cDNA from 2 µg of total RNA. The primer sequences used in this study are listed in Supplementary Table S1. The primers used to assess ER stress pathway–related genes were designed as described by Oslowski *et al.* (63). Real-time PCR amplification was performed using a CFX Connect Real-Time PCR Detection System (Bio-Rad Laboratories, Singapore) with KAPA SYBR FAST qPCR Kits (KAPA Biosystems, Boston, USA). The cycling temperatures were as follows: enzyme activation at 95°C for 3 min; denaturation at 95°C for 10 s; annealing and extension at 60°C for 30 s. Data were normalized against the expression level of 18S ribosomal RNA in each sample. The experiment was performed in triplicate.

### Apoptosis detection

Apoptotic cells were detected using an Annexin V Apoptosis Detection Kit with PI (BioLegend Inc., CA, USA). HUVEC-CS cells were cultured in DMEM with or without 20 µM TP1 under normoxic or hypoxic conditions for 24 h. In some experiments, the apoptosis of HUVEC-CS cells was induced using 0.2 μg/mL doxorubicin treatment for 24 . h. After incubation, the cells were stained with FITC-conjugated annexin V and propidium iodide according to the ’manufacturer’s protocol. Apoptotic cells were quantified using the BD FACSCalibur flow cytometry system (BD Biosciences).

### ROS detection

Intracellular ROS activity was measured using a DCFDA-based fluorogenic assay. Briefly, ECs were grown as a monolayer for 24 h under various experimental conditions, washed with serum-free medium, and stained with 1 μM DCFDA for 60 min at 37°C. The dye-containing medium was then replaced with a fresh medium, and images of DCFDA-labeled cells were captured using the green fluorescence channel of the Nikon ECLIPSE Ti-S inverted microscope system. Quantification of fluorescence intensity was performed by analyzing labeled cells in ImageJ. Statistical analyses were performed using data from triplicate experiments.

### Immunostaining

Thioflavin-T staining was used to detect UPR activity. Cells were grown on coverslips under various experimental conditions for 24 h, washed with PBS, and fixed with 4% paraformaldehyde for 20 min at room temperature. The fixed cells were then permeabilized for 5 min with 0.2% Triton X-100 at room temperature, stained with 1 μg/mL DAPI for 5 min, and counterstained with 5 μM thioflavin-T for 10 min. Coverslips were then mounted on glass slides with Clarion mounting medium, and images were captured using the Nikon ECLIPSE Ti-S inverted microscope system. Quantification of fluorescence intensity was performed by analyzing labeled cells in ImageJ. The experiment was performed in triplicate.

### Aggregated protein isolation

Cells were cultured with or without TP1 under normoxic or hypoxic conditions for 24 h, after which aggregated proteins were isolated from 10 × 10^6^ cells per condition. Briefly, cells were lysed in lysis buffer [50 mM Tris–HCl (pH 7.6), 150 mM NaCl, 1 mM EDTA, 1% NP-40, 0.1% SDS, 0.5% SDC, and protease inhibitor cocktail] with sonication at 4°C. Cell debris was removed via centrifugation at 3000 × *g*. Aggregated proteins were isolated from precleared lysates via ultracentrifugation at 112,000 × *g* and 4°C for 1 h. Aggregates were washed once with lysis buffer and dissolved in a modified lysis buffer containing 5% ammonium hydroxide. Bradford quantification was used to measure the protein content. The experiment was performed in triplicate.

## Supplementary information

Supplementary Data 1 (PDF); Supplementary Data 2 (Excel files) and Supplementary Data 3 (Excel files).

## Availability of data and material

The mass spectrometry proteomics raw data files, along with the Proteome Discoverer search data file (including protein summary), have been deposited to the ProteomeXchange Consortium (http://proteomecentral.proteomexchange.org) via the jPOSTrepo (Japan ProteOme STandard Repository) partner repository (64) with the dataset identifier PXD039938 for ProteomeXchange and JPST001996 for jPOST.

## Author contributions

BD. involved in designing and performing experiments, analyzing data, and writing the manuscript. SL and AK performed peptide extraction, purification, and confocal microscopic-based cellular uptake studies. SKS conceived and oversaw the proteomics analysis and edited the manuscript. JPT conceived the idea, supervised, and edited the manuscript. All authors have read and approved the manuscript.

## Competing interests

The authors declare that they have no potential conflict of interest.

## Acknowledgments

This study was supported in part by the Nanyang Technological University Internal 702 Funding to Synzymes and Natural Products (SYNC) and the AcRF Tier 3 funding (MOE2016703 T3-1-003).

^1^

## Footnotes

1 CRP = Cysteine-rich peptides; DMEM = Dulbecco’s modified Eagle medium; EC = Endothelial cells; ER = Endoplasmic reticulum; FBS = Fetal bovine serum; PBS = Phosphate-buffered saline; NO= Nitric oxide; ROS = Reactive oxygen species; SD = Standard deviations; SEM = Standard error of the mean; UPR = Unfolded protein response; VED = Vascular endothelial dysfunction

## Notes

### Competing Interest Statement

The authors have declared no competing interest.

