## Supplementary material for "Ginsentide TP1 Protects Hypoxia-Induced Dysfunction and ER Stress-Linked Apoptosis": Supplimentroy figures and tables

Running title: TP1 prevents hypoxia-induced endothelial dysfunction

***Correspondence:**
Prof James P Tam, PhD
School of Biological Sciences
Synthetic Enzymes and Natural Products Center
Nanyang Technological University,
60 Nanyang drive, Singapore 637551
Tel: (+65) 6316-2833


**Supplementary Figures**

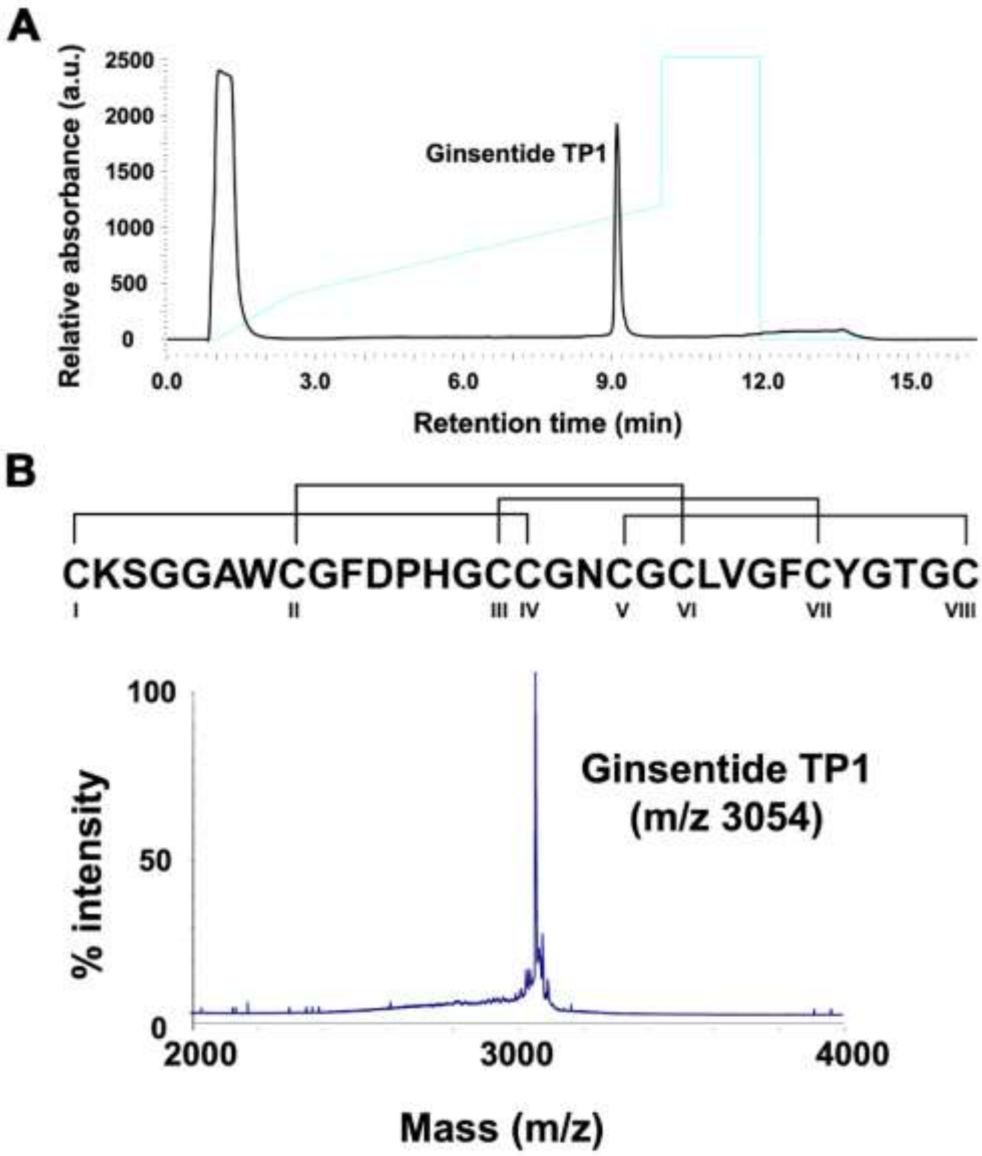


**Fig. S1: HPLC and mass spectrometry profiles of purified ginsentide TP1 extracted from *Panax ginseng* flowers.** A) HPLC chromatogram of purified TP1. B) Mass spectrometry profile of purified TP1.

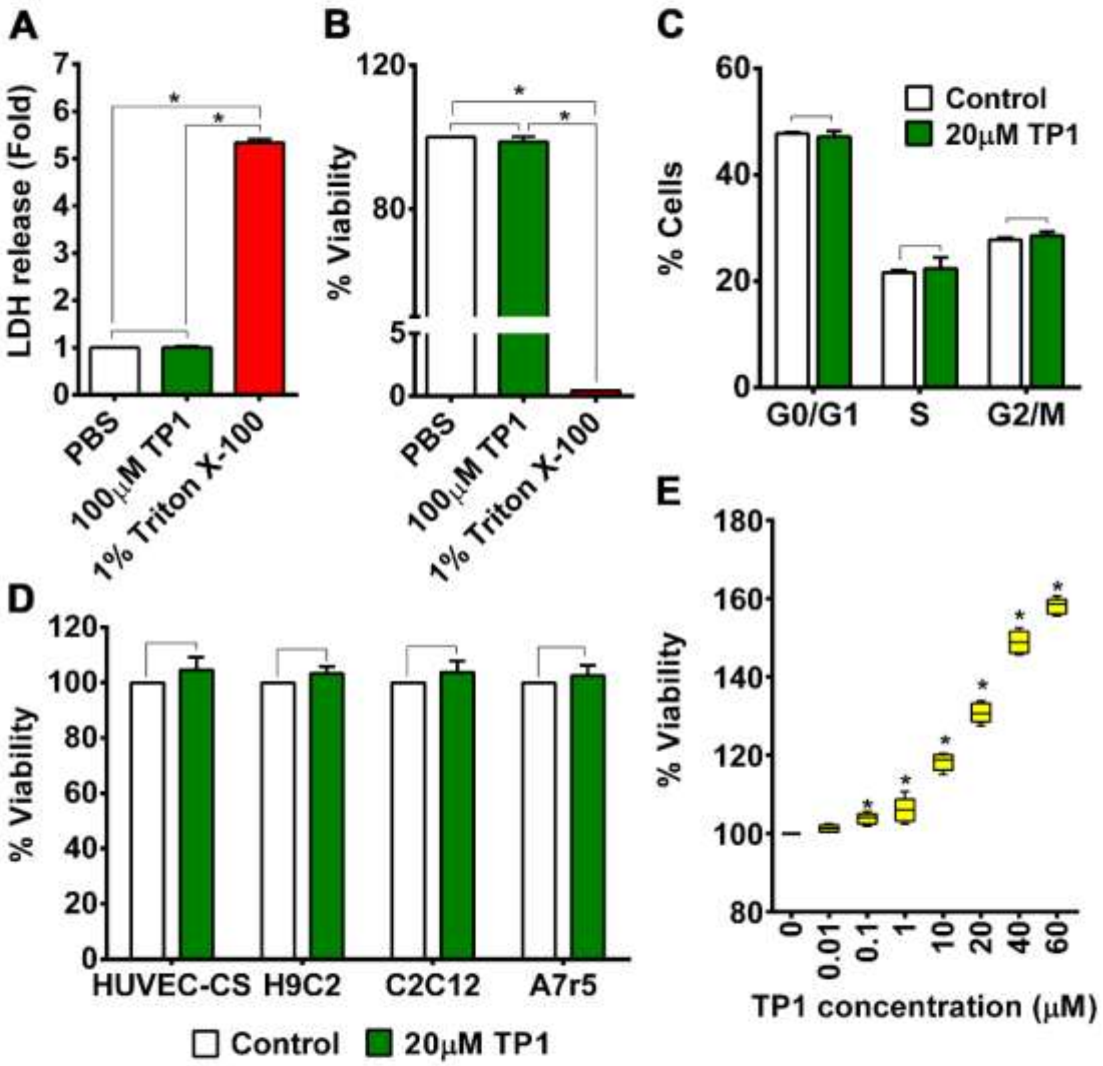


**Fig. S2: Toxicity and dose–response curve of ginsentide TP1.** A, B) Effect of TP1 on cell viability/cytotoxicity and membrane integrity in HUVEC-CS cells. Precultured cells were treated with 100 μM TP1 for 24 h. PBS and 1% TritonX-100 were used as a vehicle, and positive (cell death) controls, respectively. Cytotoxicity and membrane-damaging effects were measured using an LDH release assay (A), and cell viability was assessed using an MTT assay (B). Relative quantifications are expressed as means ± standard deviations (SDs), and statistical significance was calculated using four biological replicates. *$P < 0.05$. C) Cell cycle analysis of 20 μM TP1–treated and vehicle control–treated HUVEC-SC cells. DNA content of the cells was measured using flow cytometry after propidium iodide staining. Data from cells in different cell cycle phases are expressed as means ± SDs, and statistics were calculated using triplicate biological replicates. D) Viability of different cells was measured using an MTT assay after treatment with 20 μM TP1 or vehicle control. Relative survival is expressed as mean values, and statistical significance was calculated using three independent experimental replicates. E) Dose–response curve of TP1. HUVEC-CS cells were cultured and treated with different doses (0–60μM) of TP1 for 24 h in a hypoxic environment, and an MTT assay was used to assess cell viability. Data are means ± standard error of the mean of five independent experimental replicates. *$P < 0.05$.

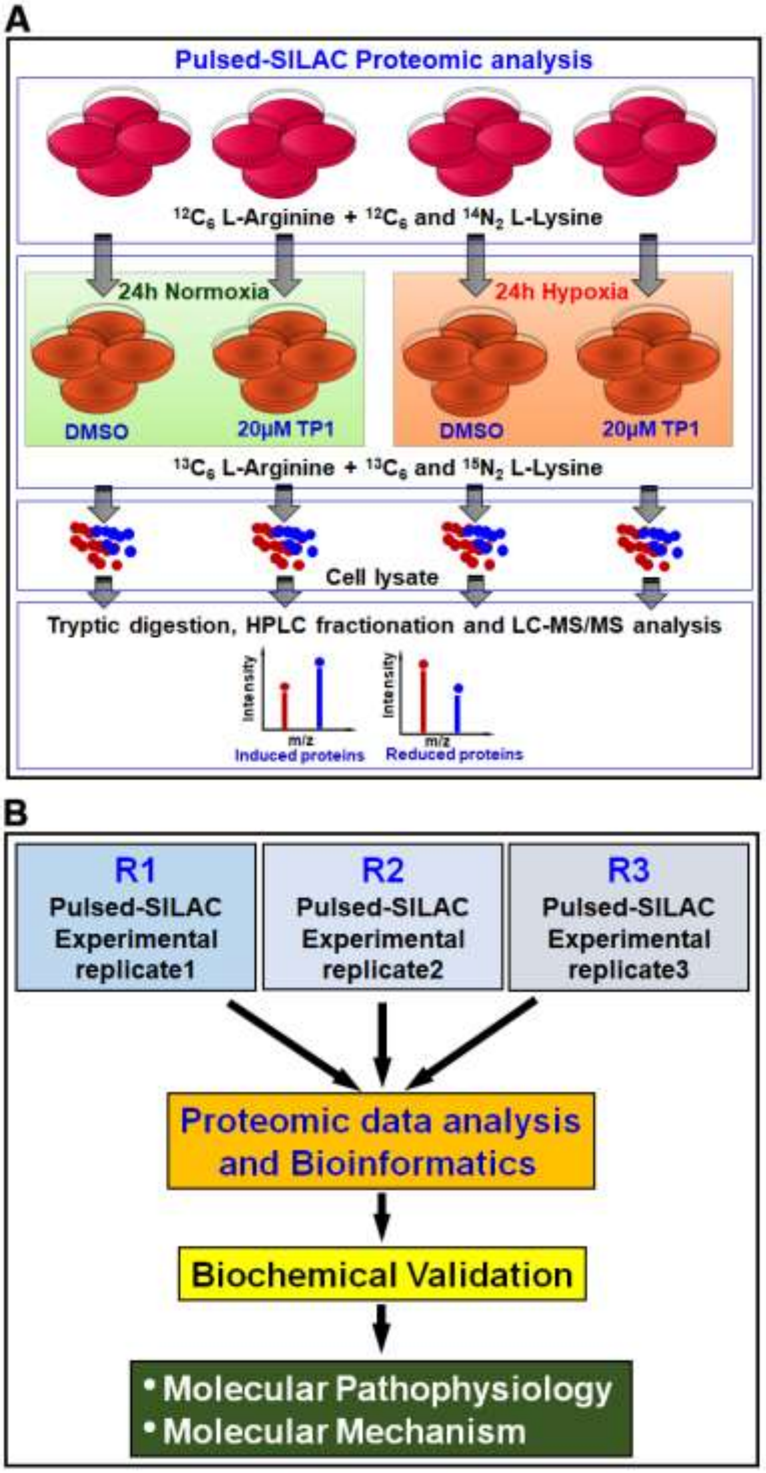


**Fig. S3: Schematic representation of the experimental workflow of pulsed SILAC labeling and proteomics.** A) Schematic representation of the pulsed SILAC–based proteomics experiment. B) Overall experimental workflow. R1, R2, and R3 represent triplicate experiments.

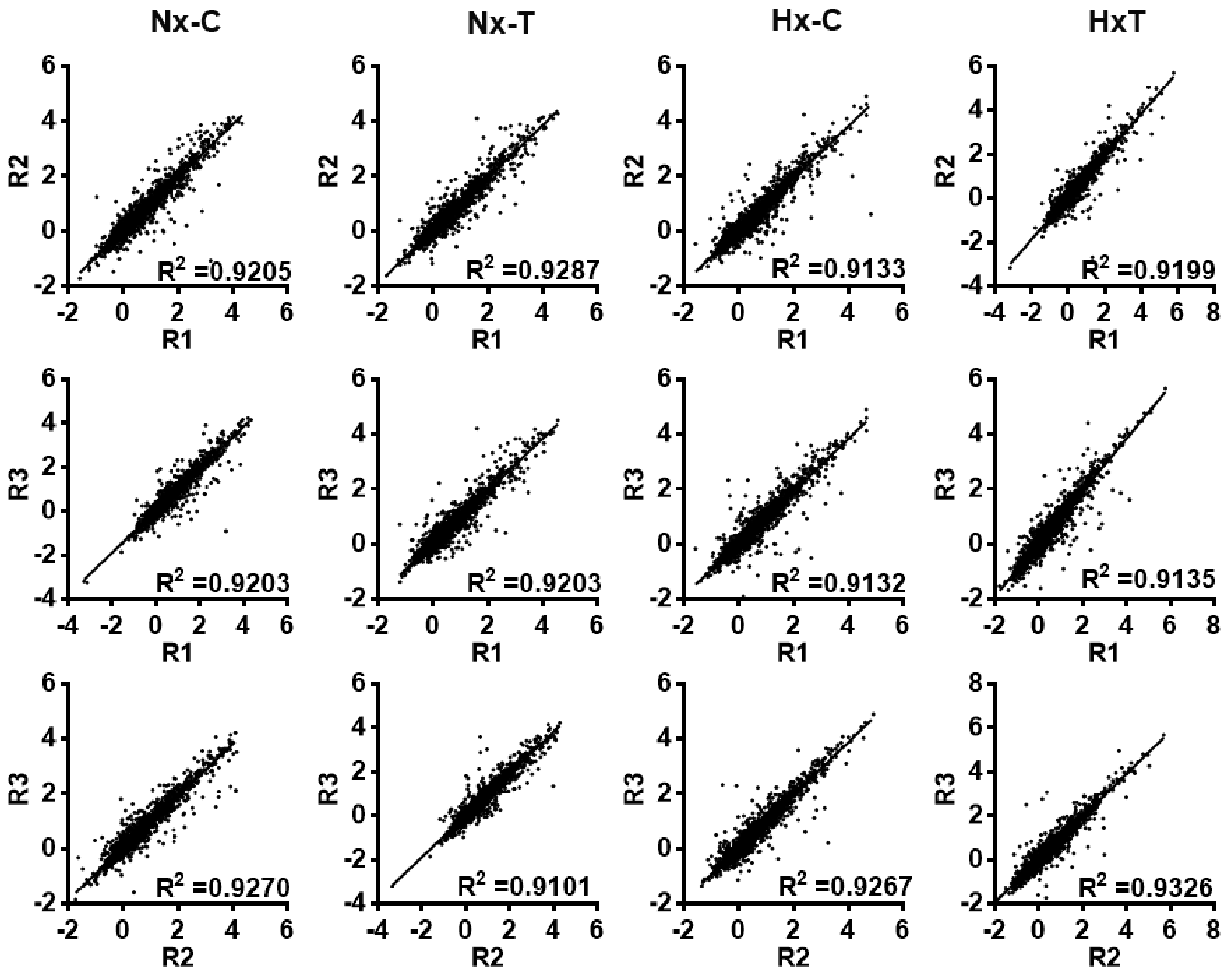


**Fig. S4: Linear regression plots representing the correlation among triplicate experiments under different experimental conditions**. X and Y axes represent the $log_2$-transformed heavy/light (H/L) SILAC ratio of proteins from respective experimental replicates. $R^2$ value represents the correlation between replicates, with the high correlations indicating the reliability of the detection of the pulsed SILAC experiment.

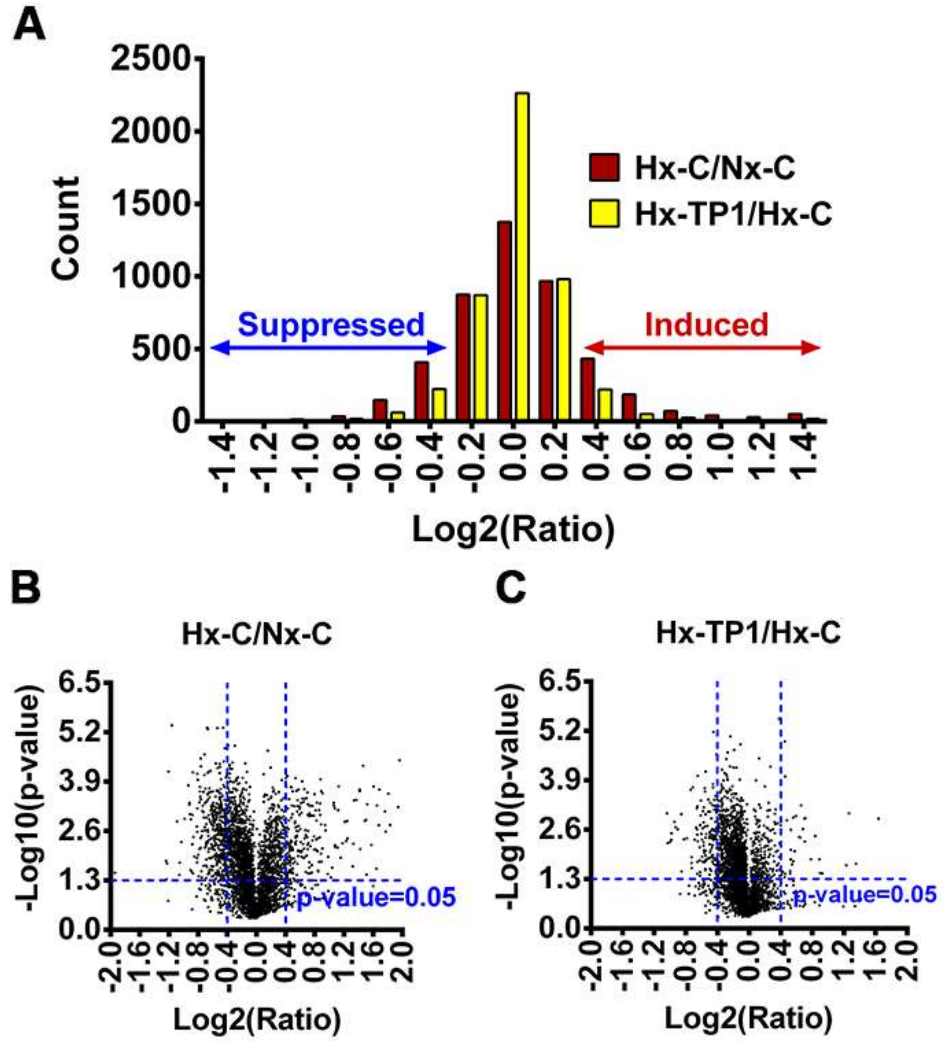


**Fig. S5: Statistical analysis of proteomic data.** A) Frequency distribution plot of fold-change ratio. Cutoff ratio values of <0.76 [log2 (Ratio) < −0.4] and >1.32 [log2 (Ratio) > 0.4] were set for the suppressed and induced proteins, respectively. B, C) Volcano plot highlighting statistically significant differential protein synthesis during hypoxia (ratio of Hx-C/Nx-C) (B) and TP1 treatment during hypoxia (Hx-TP1/Hx-C) (C). HUVEC-CS cells were cultured, SILAC-labeled, and quantified under the respective experimental conditions, including normoxic (Nx-C)/hypoxic (Hx-C) vehicle control and normoxic (Nx-TP1)/hypoxic (Hx-TP1) TP1-treated conditions. Statistical analysis was performed using triplicate experiments. *P < 0.05.

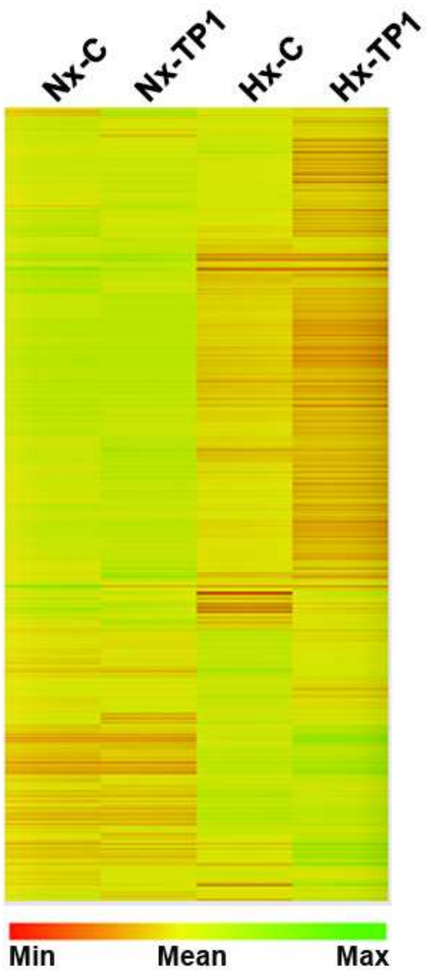


**Fig. S6: Differential abundance of newly expressed proteins in HUVEC-CS cells with or without TP1 treatment under normoxic and hypoxic conditions.** Heat map representing a global view of the differentially synthesized new proteins under the indicated experimental conditions. HUVEC-CS cells were treated with 20 µM TP1 for 24 h under normoxic or hypoxic conditions, and PBS was used as vehicle control. A SILAC-based protein labeling approach was used to label the newly synthesized proteins, which were later quantified using an LC−MS/MS-based quantitative proteomic technique. Cluster analysis of the identified proteins was performed using the online bioinformatics tool Gene Pattern (http://genepattern.broadinstitute.org) with hierarchical clustering and Pearson correlation options. Vehicle control under normoxic (Nx-C) or hypoxic (Hx-C) conditions, and TP1 treatment under normoxic (Nx-TP1) or hypoxic (Hx-TP1) conditions.

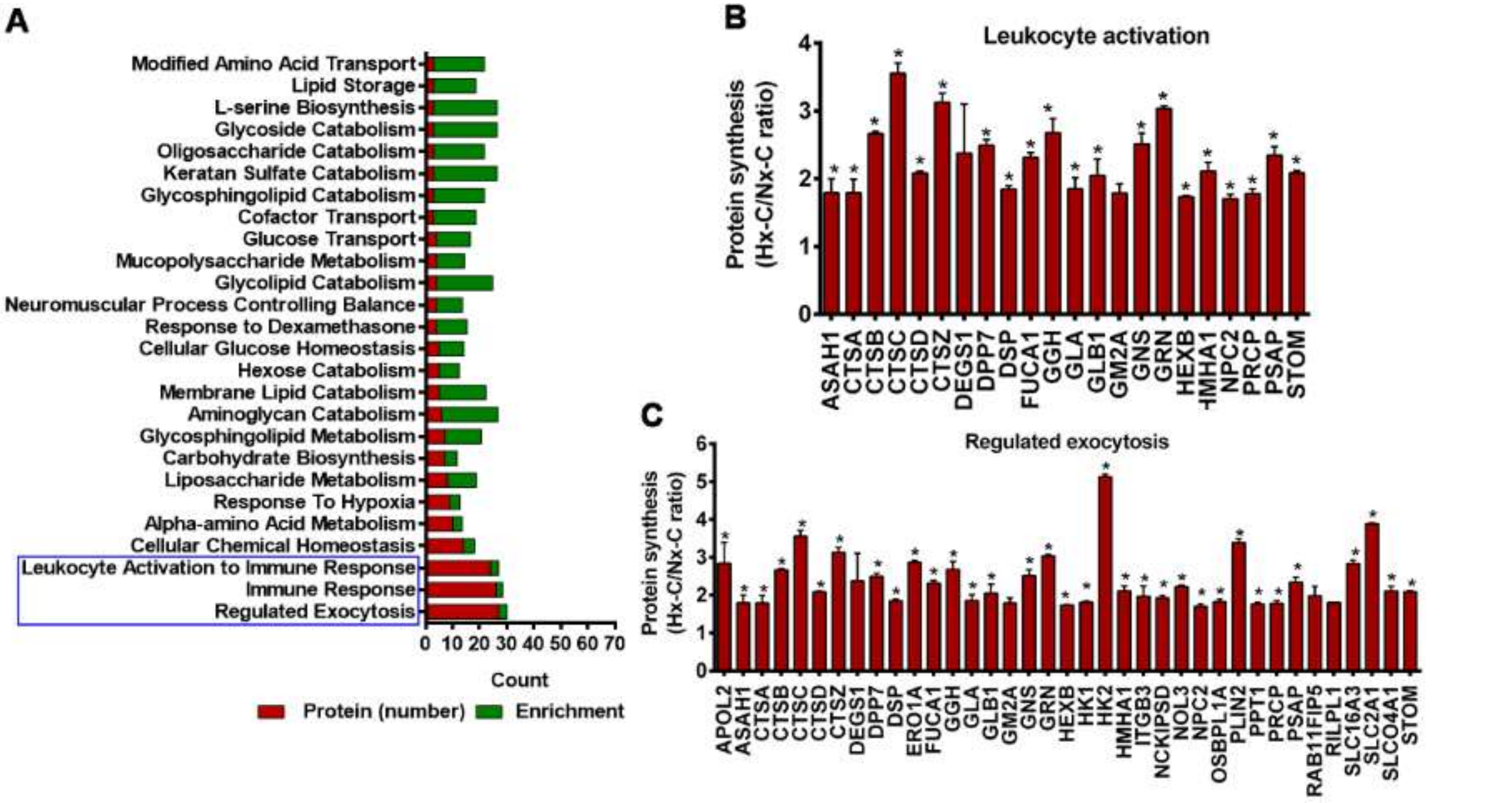


**Fig. S7: Hypoxia-induced biological processes in endothelial cells (ECs).** A) Bioinformatics-based enrichment analysis of the hypoxia-induced biological processes in ECs. B, C) Relative abundance of hypoxia-induced proteins involved in the regulation of bioprocesses, including immune responses and leukocyte activation (B) and exocytosis (C). Quantitation values are means ± standard error of the mean calculated from triplicate experiments. *$P < 0.05$.

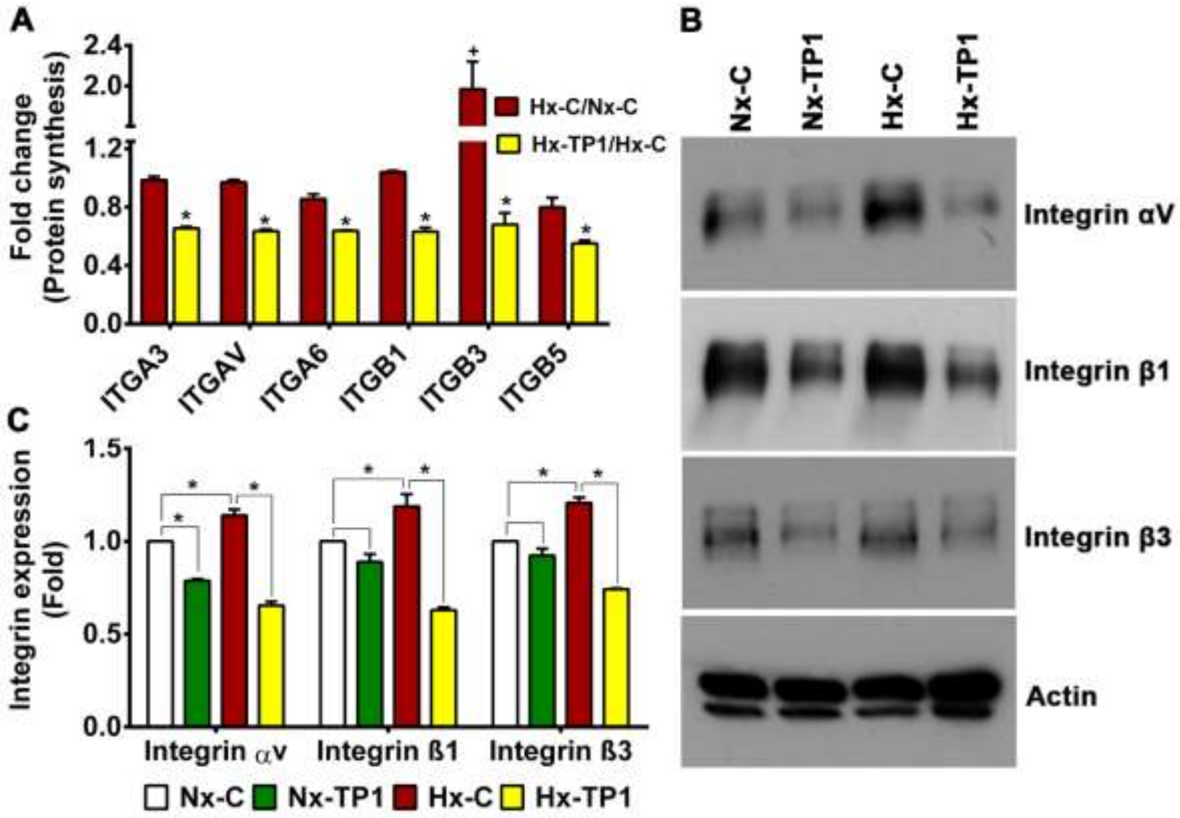


**Fig. S8: Ginsentide TP1 suppresses integrin expression in hypoxic endothelial cells.** A) Relative change in newly synthesized integrin subtypes under the hypoxic condition (Hx-C/Nx-C) and TP1 treatment under the hypoxic condition (Hx-TP1/Hx-C). Significance $^{+}P < 0.05$ vs. Nx-C and $*P < 0.05$ vs. Hx-C. B) Western blot images showing the expression of integrin subtypes under the indicated experimental conditions. C) Graphical representation of the normalized expression of integrin subtypes under the respective experimental conditions. HUVEC-CS cells were cultured and treated with PBS or 20 µM TP1 under normoxic or hypoxic conditions for 24 h. Quantitation values are means ± standard error of the mean, which were calculated using triplicate experiments. $*P < 0.05$.

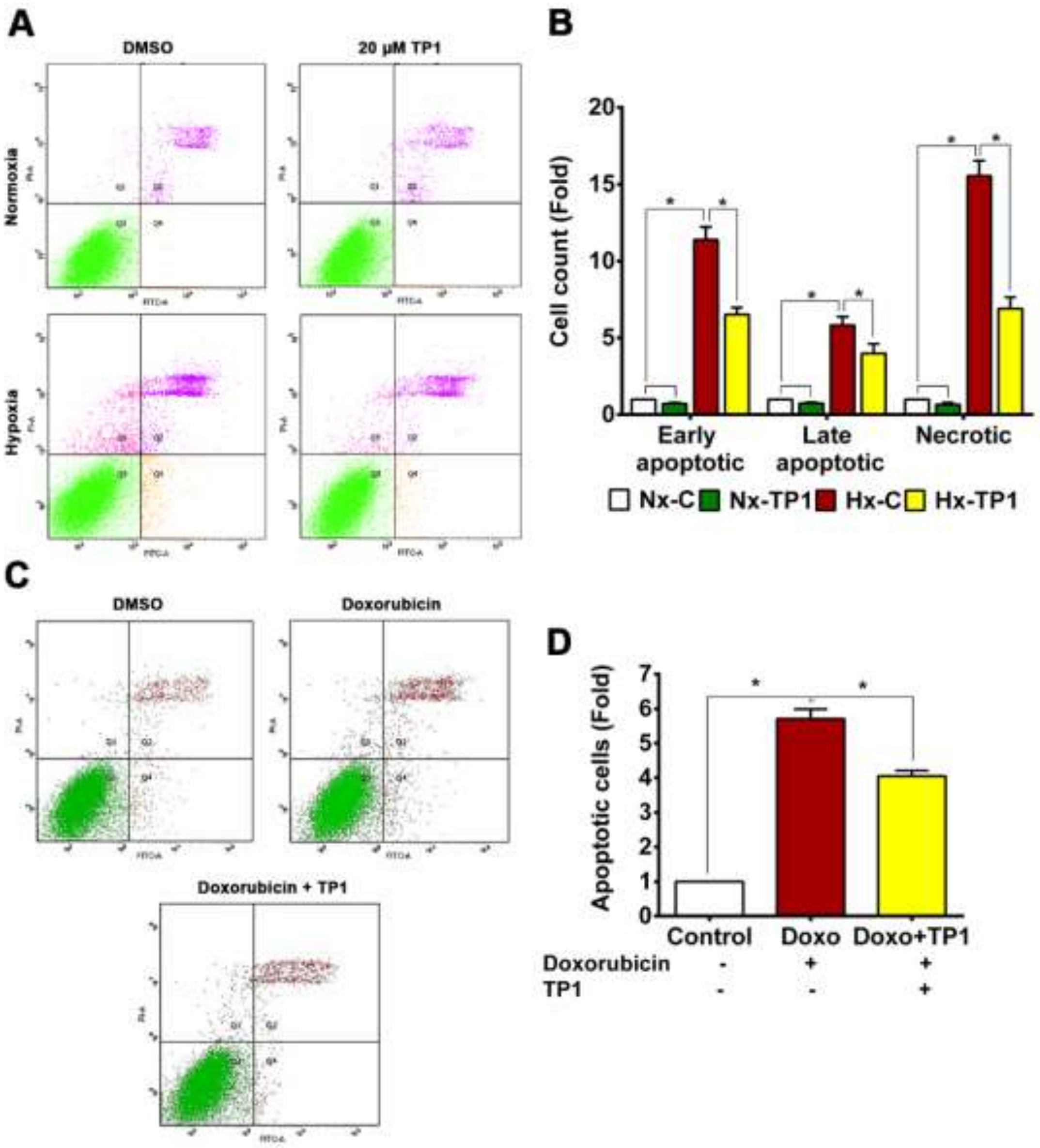


**Figure S9: TP1 prevents apoptosis in HUVEC-CS.** A) Annexin V and PI staining in HUVEC-CS cells treated with 20 µM TP1 and DMSO at normoxic (Nx-C) or hypoxic (Hx-C) conditions. The third quadrant represents living cells (Annexin V, PI negative), the first one early apoptotic cells (Annexin V positive, PI negative), the second late apoptotic (Annexin and PI positive), and the fourth necrotic or dead cells (Annexin V negative and PI positive). B) Relative quantification was based on cell count. Statistical significance was calculated using three independent experimental replicates. Data are means ± standard error of the mean for triplicate experiments. *P < 0.05. D) Annexin V and PI staining of HUVEC-CS cells treated with 0.2µg/ml doxorubicin with or with 20 µM TP1 co-treatment. DMSO was used as vehicle control. E) Relative quantification of doxorubicin-induced apoptotic cells under respective treatment conditions. Data are means ± standard error of the mean for triplicate experiments. *P < 0.05.

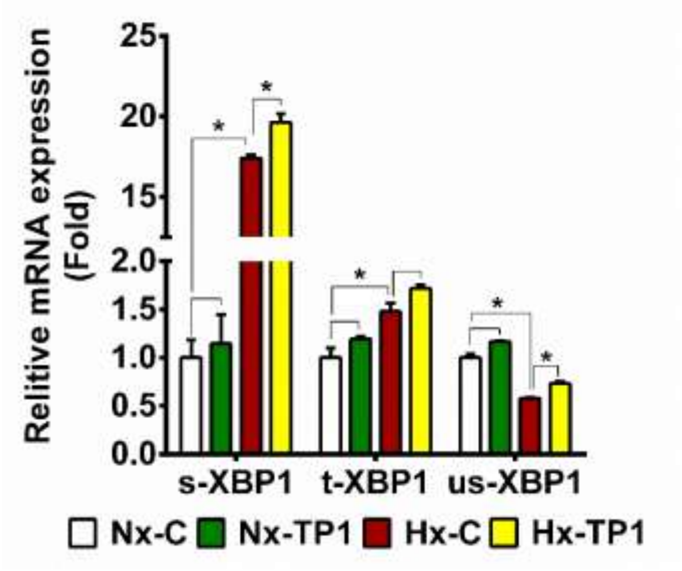


**Fig. S10: Effect of TP1 on the hypoxia-induced unfolded protein response (UPR) in endothelial cells.** Relative mRNA expression levels of the UPR marker gene XBP1, including total XBP (t-XBP1), its spliced isoform (s-XBP1), and its unspliced isoform (us-XBP1), under the indicated experimental conditions. Quantitation values are means with ± standard deviations from triplicate biological replicates. *$P < 0.05$. HUVEC-CS cells were cultured and treated with PBS or TP1 for 24 h under normoxic or hypoxic conditions. Vehicle control normoxic (Nx-C) and hypoxic (Hx-C) and TP1-treated normoxic (Nx-TP1) and hypoxic (Hx-TP1) treatments are shown.

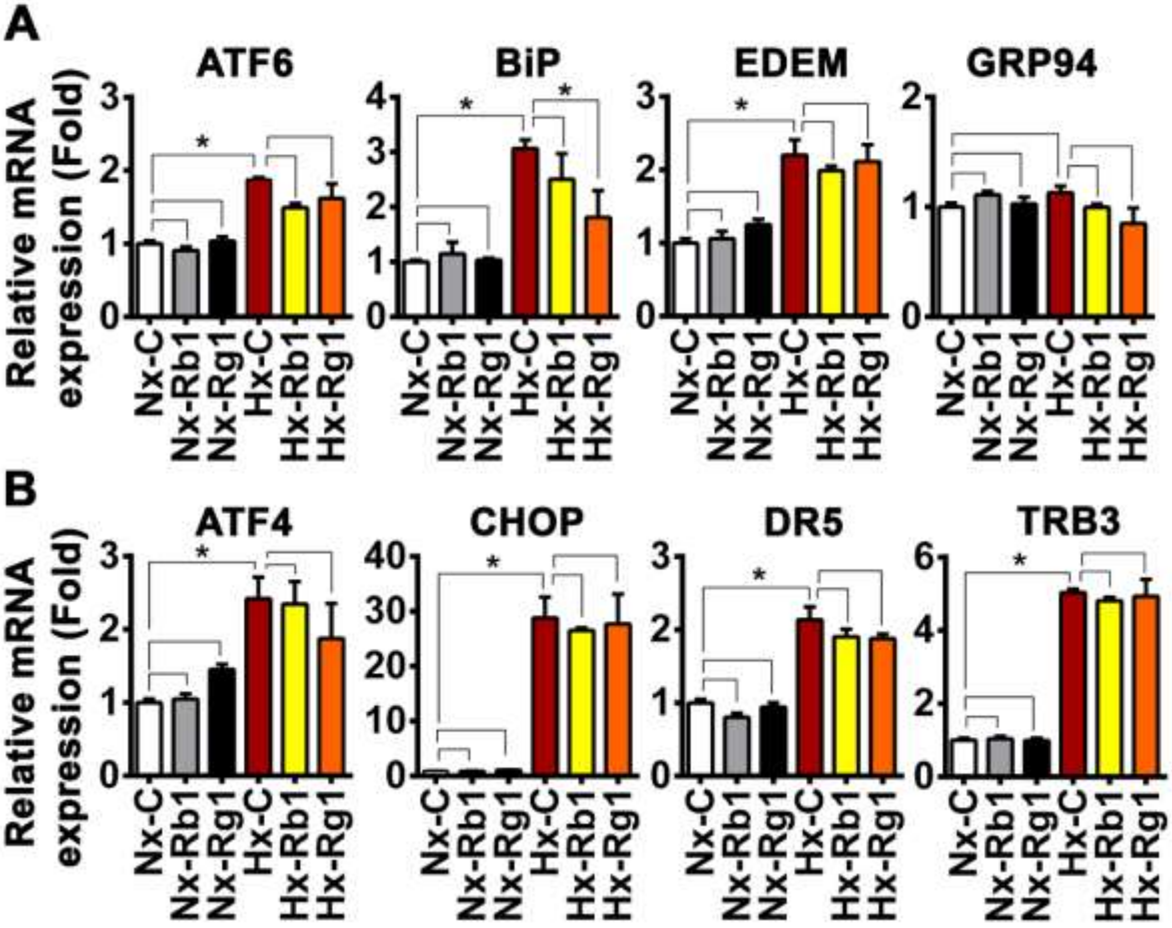


**Fig. S11: Effects of ginsenosides on hypoxia-induced ER stress–related gene expression in ECs.** A, B) Relative mRNA expression levels of ER stress–related adaptive unfolded protein response (UPR) transduction cascade genes (A) and UPR-mediated death signaling cascade genes (B) in HUVEC-CS cells cultured under the indicated experimental conditions. Means ± standard error of the mean were calculated from triplicate biological replicates. Vehicle control cells were cultured with PBS for 24 h under normoxic (Nx-C) or hypoxic (Hx-C) conditions, and 20 µM ginsenoside Rb1- or Rg1-treated cells were cultured for 24 h under normoxic (Nx-Rb1/Nx-Rg1) or hypoxic (Hx-Rb1/Hx-Rg1) conditions. *P < 0.05.

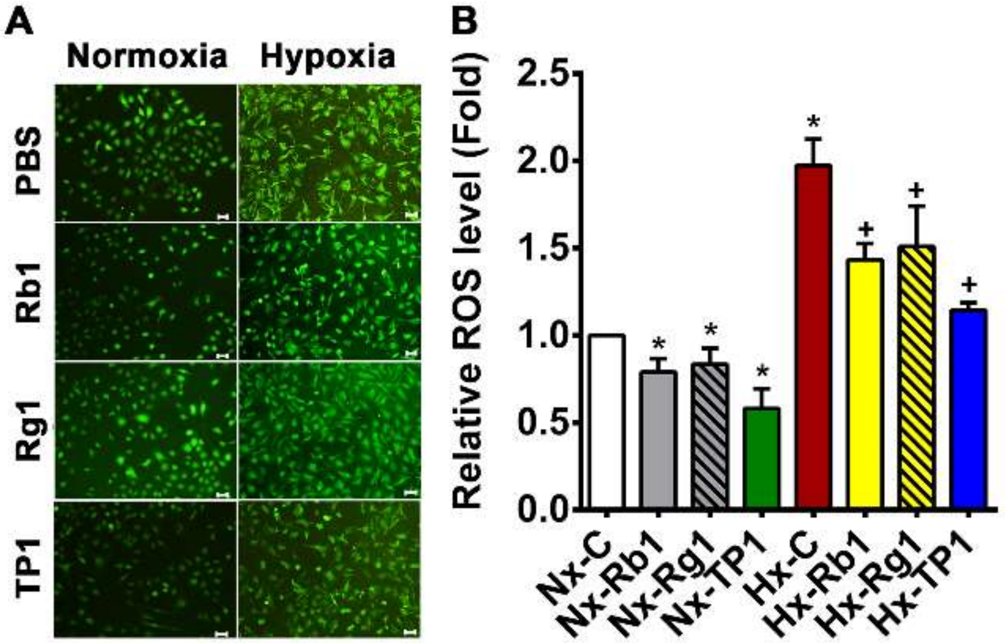


**Fig. S12: Effect of ginsenoside treatment on the intracellular ROS load in ECs.** A) Effect of Rb1 and Rg1 on intracellular ROS levels in HUVEC-CS cells cultured under the indicated experimental conditions. Scale bar: 25 µm. B) Relative quantification was performed using relative fluorescence intensity, and statistical significance was calculated using three biological replicates. Data are means ± standard error or the mean. Normoxic or hypoxic HUVEC-CS cells were treated with PBS (vehicle control), 20 µm ginsenoside Rb1 or Rg1, or 20 µm ginsentide TP1 (positive control) for 24 h. *P < 0.05.

**Full blot images.** Western blot analysis of HUVEC-CS endothelial cells on 20µM ginsentide TP1 treatment under normoxic and hypoxic conditions. Vehicle control cells were cultured for 24h under normoxia (Nx-C) or hypoxia (Hx-C) conditions. Cells were treated with TP1 for 24h under normoxia (Nx-TP1) or hypoxia (Hx-TP1)

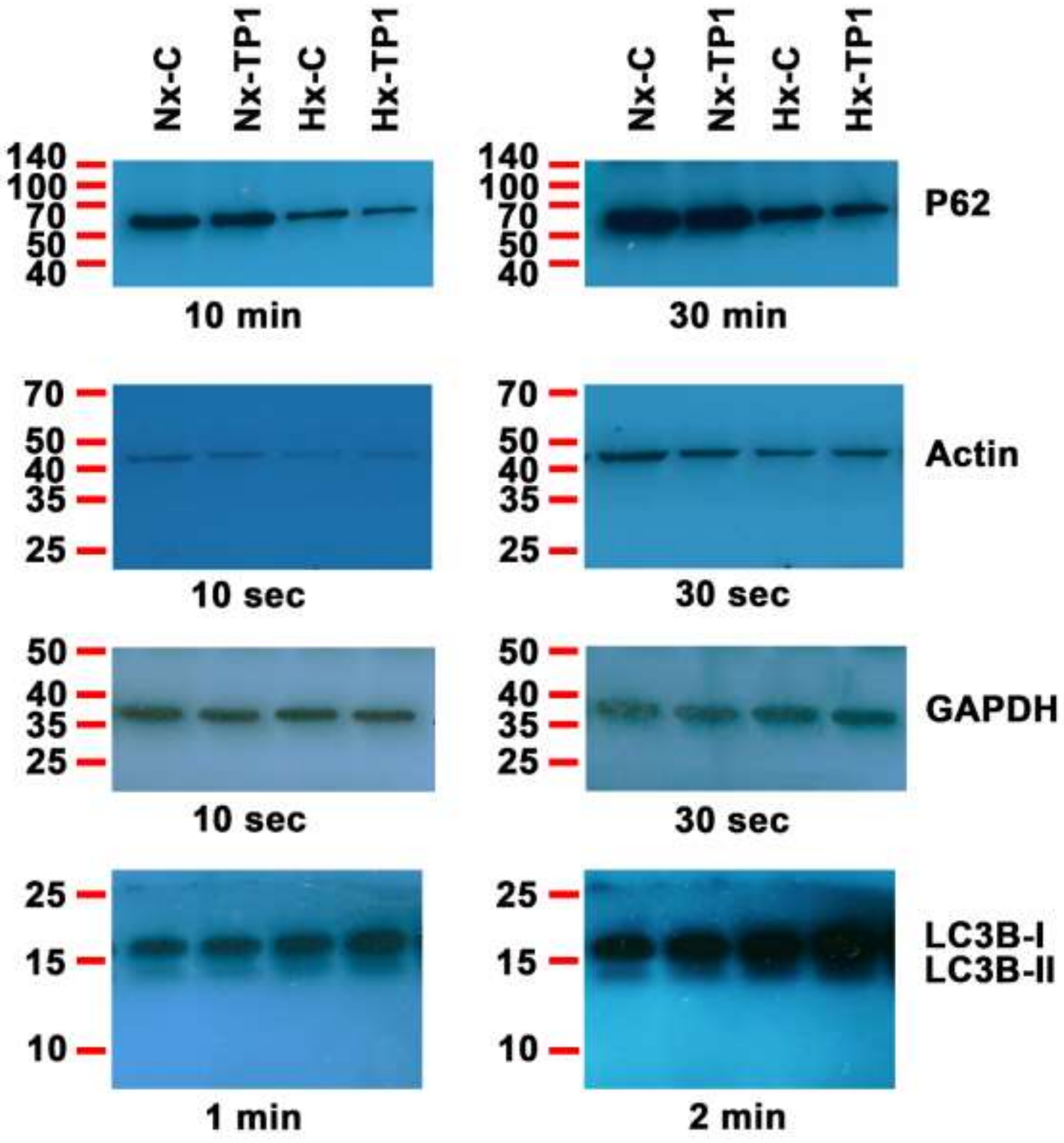

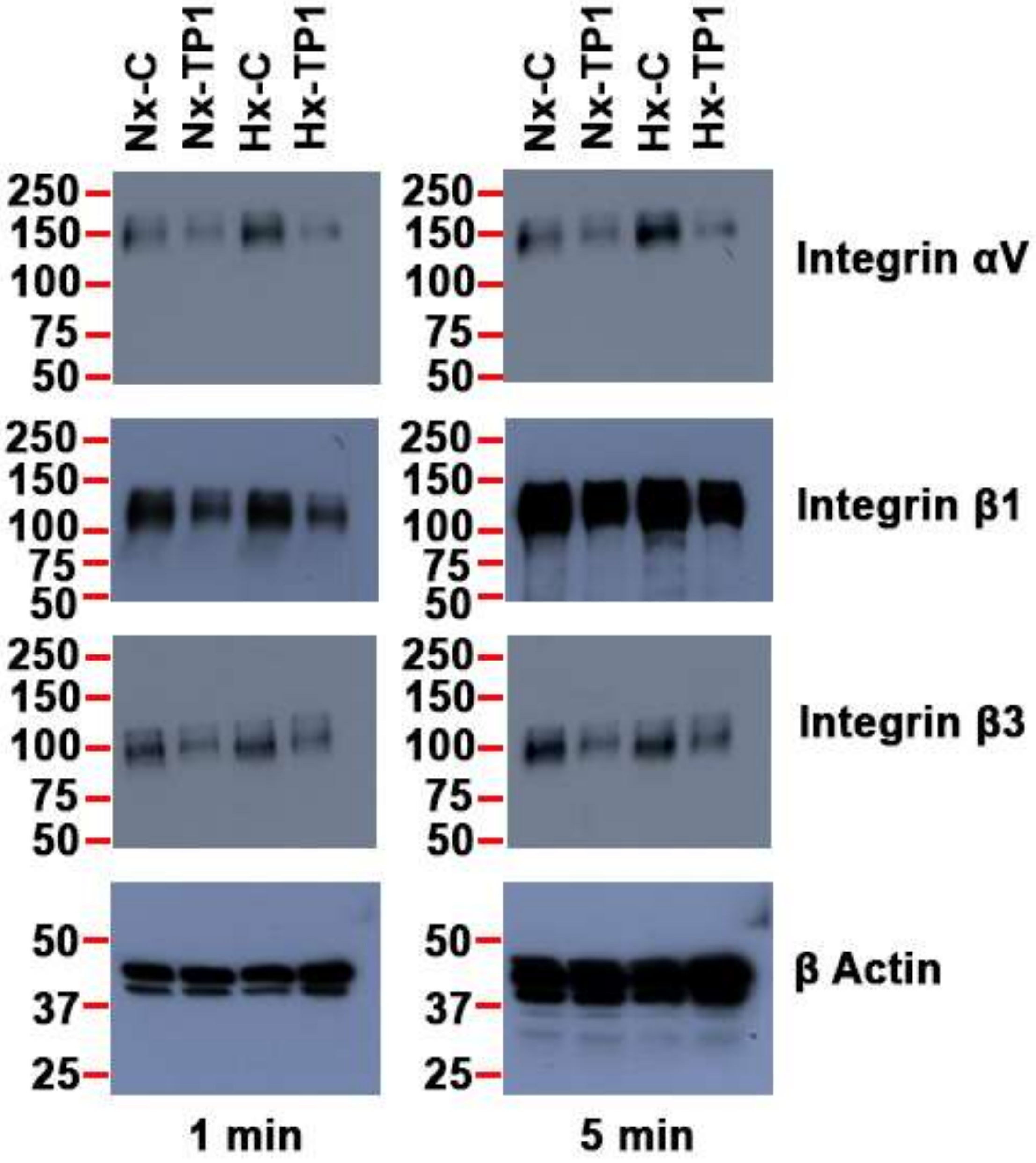
Nx-C
Nx-TP1
Hx-C
Hx-TP1
250
150
100
75
50
37
25
Integrin αV
Integrin β1
Integrin β3
β Actin
1 min
5 min

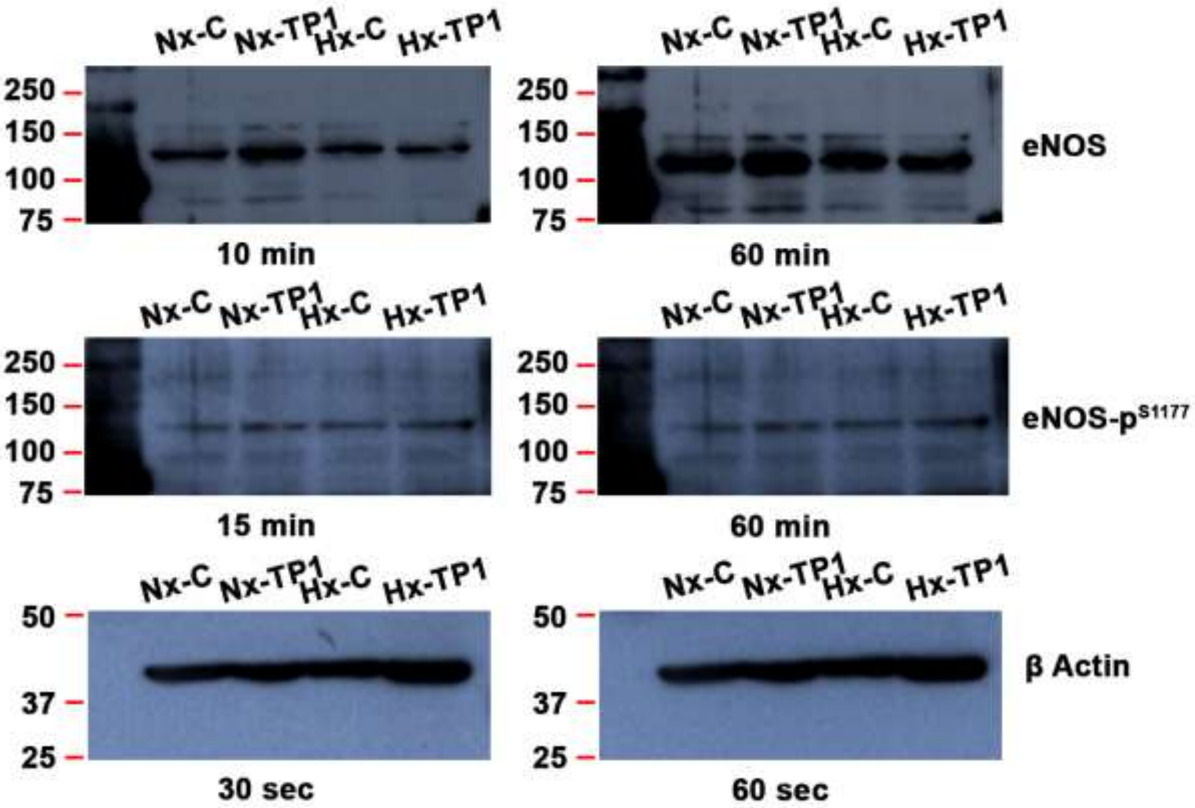
Nx-C
Nx-TP1
Hx-C
Hx-TP1
250
150
100
75
eNOS
10 min
60 min
15 min
eNOS-p$^{S1177}$
β Actin
50
37
25
30 sec
60 sec

### Supplementary Tables

**Table S1: List of genes and corresponding primers used in quantitative RT-PCR.**

| Genes | Forward primer sequence (5´–3´) | Reverse primer sequence (5´–3´) |
|---|---|---|
| **ALCAM** | CGCAATGCAACAGGAGACTA | GGCTAGATCGAAGCCTGATG |
| **ICAM1** | GGCTGGAGCTGTTTGAGAAC | ACTGTGGGGTTCAACCTCTG |
| **L1CAM** | GCCAAAGGAGACAGTGAAGC | GCGTGGCAGATGTAGTCTGA |
| **VCAM1** | CAGACAGGAAGTCCCTGGAA | TTCTTGCAGCTTTGTGGATG |
| **ATF4** | GTTCTCCAGCGACAAGGCTA | ATCCTGCTTGCTGTTGTTGG |
| **CHOP** | AGAACCAGGAAACGGAAACAGA | TCTCCTTCATGCGCTGCTTT |
| **DR5** | CACCAGGTGTGATTCAGGTG | CCCCACTGTGCTTTGTACCT |
| **TRB3** | TGGTACCCAGCTCCTCTACG | TTCTCCAGCACCAGCTTCTT |
| **ATF6** | GCCTTTATTGCTTCCAGCAG | TGAGACAGCAAAACCGTCTG |
| **BiP** | TGTTCAACCAATTATCAGCAAACT | TTCTGCTGTATCCTCTTCACCAGT |
| **EDEM** | CAAGTGTGGGTACGCCACG | AAAGAAGCTCTCCATCCGGTC |
| **GRP94** | GAAACGGATGCCTGGTGG | GCCCCTTCTTCCTGGGTC |
| **sXBP1** | CTGAGTCCGAATCAGGTGCAG | ATCCATGGGGAGATGTTCTGG |
| **usXBP1** | CAGCACTCAGACTACGTGCA | ATCCATGGGGAGATGTTCTGG |
| **Total-XBP1** | TGGCCGGGTCTGCTGAGTCCG | ATCCATGGGGAGATGTTCTGG |
| **RPLP0** | TCGACAATGGCAGCATCTAC | GCCTTGACCTTTTCAGCAAG |
| **RPL13A** | GTACGCTGTGAAGGCATCAA | CGCTTTTTCTTGTCGTAGGG |
| **B-actin** | AGAGCTACGAGCTGCCTGAC | AGCACTGTGTTGGCGTACAG |
| **18S** | GTAACCCGTTGAACCCCATT | CCATCCAATCGGTAGTAGCG |

sXBP1: Spliced XBP1 and usXBP1: Unspliced XBP1
